# A gut-brain-gut interoceptive circuit loop gates sugar ingestion in *Drosophila*

**DOI:** 10.1101/2024.09.02.610892

**Authors:** Xinyue Cui, Matthew R. Meiselman, Staci N. Thornton, Nilay Yapici

## Abstract

The communication between the brain and digestive tract is critical for optimising nutrient preference and food intake, yet the underlying neural mechanisms remain poorly understood^1–7^. Here, we show that a gut-brain-gut circuit loop gates sugar ingestion in flies. We discovered that brain neurons regulating food ingestion, IN1^8^, receive excitatory input from enteric sensory neurons, which innervate the oesophagus and express the sugar receptor *Gr43a*. These enteric sensory neurons monitor the sugar content of food within the oesophagus during ingestion and send positive feedback signals to IN1s, stimulating the consumption of high-sugar foods. Connectome analyses reveal that IN1s form a core ingestion circuit. This interoceptive circuit receives synaptic input from enteric afferents and provides synaptic output to enteric motor neurons, which modulate the activity of muscles at the entry segments of the crop, a stomach-like food storage organ. While IN1s are persistently activated upon ingestion of sugar-rich foods, enteric motor neurons are continuously inhibited, causing the crop muscles to relax and enabling flies to consume large volumes of sugar. Our findings reveal a key interoceptive mechanism that underlies the rapid sensory monitoring and motor control of sugar ingestion within the digestive tract, optimising the diet of flies across varying metabolic states.

## Main

Most animals regulate food intake via a complex network of sensory, homeostatic, and hedonic physiological processes that are under the control of neural circuits within the central and enteric nervous systems. Communication between the brain and gastrointestinal tract is essential for assessing the body’s metabolic status, regulating the ingestion of specific nutrients, and triggering satiety responses when energy reserves are replenished^1–3, 6^. Recent studies highlight the critical roles of gut-brain circuits in regulating homeostatic and hedonic aspects of food intake across species, including invertebrates^9–13^ and mammals^14–21^. In mammals, the vagus nerve serves as one of the primary pathways for bidirectional communication between the gastrointestinal tract and the brain. As part of the parasympathetic nervous system, it extensively innervates multiple compartments of the digestive tract, including the oesophagus, stomach, small intestine, and parts of the large intestine^2, 6^. This broad network of innervation allows the vagus nerve to play critical roles in regulating food ingestion, nutrient preference and various digestive processes, such as swallowing, gastric secretions, and gut motility^2, 6, 16, 18–24^. Despite the critical roles of the gut-brain axis in regulating food intake and metabolism, the interoceptive circuits that translate sensory signals from the gut into motor actions that control nutrient preference and ingestion remain poorly understood. It remains a challenge to address this knowledge gap in mammals due to the complexity of their enteric nervous system. The small enteric nervous system of *Drosophila* shares many functions with its vertebrate counterparts^3, 4, 7^, providing a valuable model for studying the functional principles of gut-brain circuits.

The fly digestive tract is innervated by neurons originating from the stomatogastric nervous system, which encompasses the hypocerebral ganglion (HCG) and central neurons located in the brain and the ventral nerve cord^3, 5, 13^. Neurons with cell bodies located in the pars intercerebralis (PI) region of the brain and HCG innervate the crop and the anterior regions of the midgut. Meanwhile, neurons in the abdominal ganglia of the ventral nerve cord extend their arborisations to the posterior regions of the midgut, as well as to the hindgut^3, 12^. Recent studies have revealed that the PI and enteric sensory neurons expressing the mechanosensory channel Piezo sense crop distension and mediate food ingestion and nutrient preference in flies^9–11^. Another class of enteric neurons expressing the neuropeptide myosuppressin is remodelled by steroid hormones after mating, enabling females to consume large amounts of food^13^. These findings indicate that, as in humans and other mammals, the gut-brain axis plays significant roles in regulating nutrient preference and food ingestion in *Drosophila*.

Previously, we identified ∼12 local interneurons (IN1, for ‘‘ingestion neurons’’) in the taste processing centre of the fly brain, subesophageal zone (SEZ), as a critical regulator of food intake^8^. IN1s are persistently activated in hungry flies consuming high concentrations of sucrose but not those consuming low concentrations^8^. Furthermore, IN1 activity is essential for the rapid and precise regulation of sugar ingestion, suggesting these neurons serve as a central node in neural circuits that process taste sensory information and govern food intake^8^. Here, we use IN1s as a gateway to identify and characterise a gut-brain-gut interoceptive circuit that gates sugar ingestion across varying metabolic states.

### IN1s receive excitatory input from enteric sensory neurons expressing *Gr43a*

To identify neurons that provide sensory input to IN1s, we used a previously validated approach to map functional connectivity within the *Drosophila* nervous system^25–27^. We stimulated candidate sensory neurons with a red-shifted channelrhodopsin, Chrimson^25, 28^, while imaging the activity of IN1s using a genetically encoded calcium indicator GCaMP6s^29^ (Figs. 1a, b). Optogenetic activation of sugar-sensing neurons expressing Gr64f ^30, 31^ strongly excited IN1s. In contrast, optogenetic stimulation of other sensory neurons, such as bitter-sensing (Gr66a^32–34^), water-sensing (ppk28^35^), or mechanosensory neurons (TMC^36, 37^), did not elicit the same responses (Figs. 1c-h, Extended Data Figs. 1a-g). Next, we aimed to identify which group of sugar-sensing neurons provides sensory input to IN1s. In flies, most sugar-sensing neurons express Gr64f. These neurons are located in multiple chemosensory organs, including the labellum, legs, pharynx^30, 31, 38–42^, brain^43, 44^ and enteric nervous system^45^. We stimulated various subsets of Gr64f neurons labelled by other *Gr-GAL4s* and simultaneously recorded IN1 activity. Optogenetic stimulation of sugar-sensing neurons expressing *Gr5a*^46^, *Gr64a*^39, 40^ or *Gr64d*^38, 47^ did not elicit a significant response (Figs. 2a-c). However, stimulation of neurons expressing *Gr43a* (labelled by *Gr43a^GAL^*^4^, a knock-in insertion to *Gr43a* locus) strongly activated IN1s (Figs. 2d, g). To further narrow down the Gr43a neurons required for IN1 activation, we used two additional transgenic lines, *ChAT-GAL80*^48^ and *Gr43a-GAL4* (an enhancer-GAL4), to label subsets of Gr43a neurons. Neurons labelled by *Gr43a-GAL4* or Gr*43a^GAL^*^4^ combined with *ChAT-GAL80* were not able to activate IN1s upon optogenetic stimulation (Figs. 2e-f). The main difference among these transgenic flies was their expression in the HCG: only Gr*43a^GAL^*^4^ labelled multiple enteric sensory neurons (Figs. 2g-i, Extended Data Figs. 2a-d). Our results indicate that enteric sensory neurons expressing *Gr43a* are required for IN1 activation.

**Fig.1.**
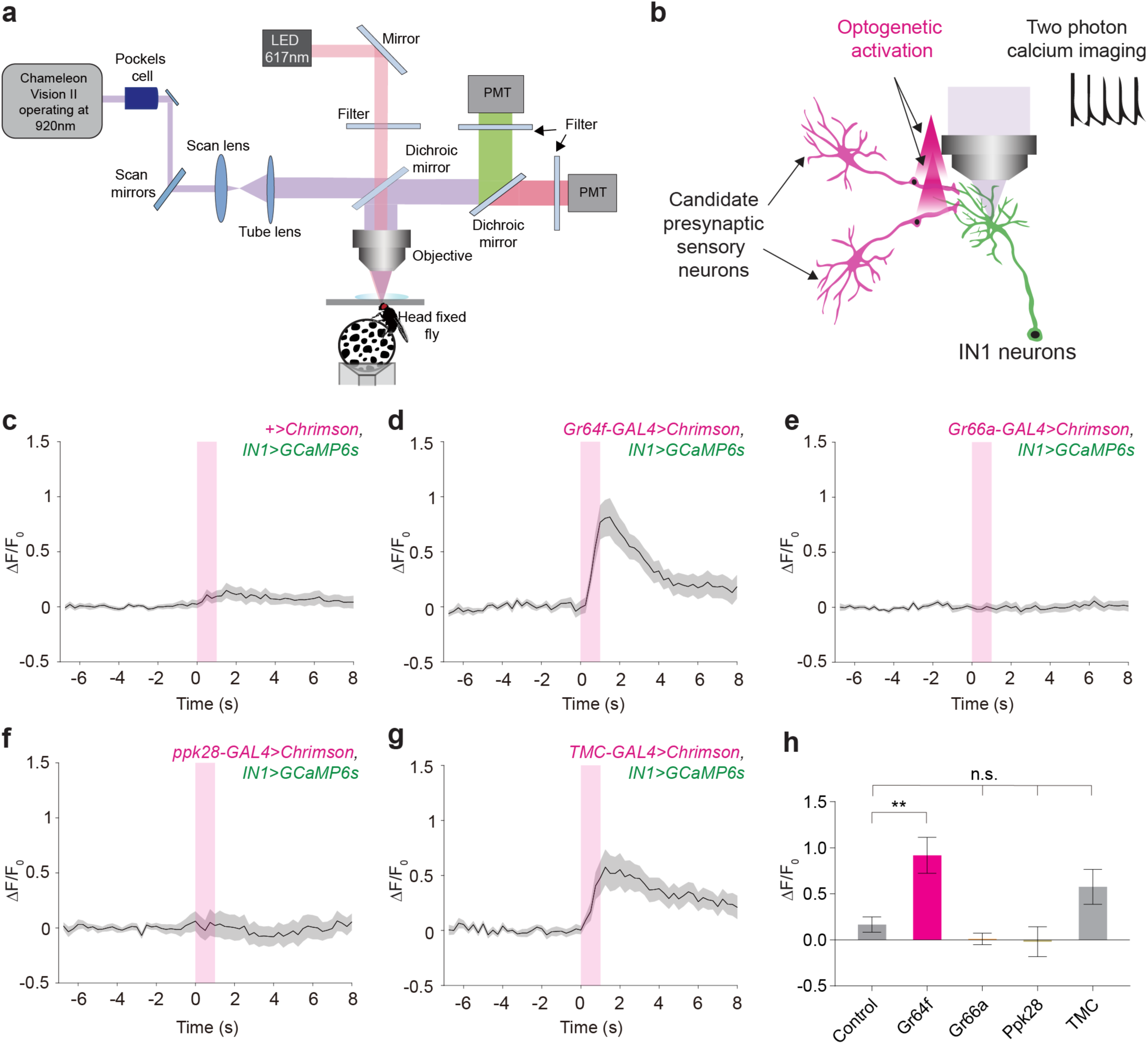
IN1s receive excitatory input from sugar-sensing neurons expressing *Gr64f*. **a**, Schematic of the two-photon microscope setup coupled with optogenetic stimulation. A male fly is standing on top of an air-suspended spherical treadmill (PMT, photomultiplier tube). **b**, Schematic showing the strategy used for optogenetic stimulation coupled with two-photon calcium imaging. GCaMP6s is expressed in IN1s, while red-shifted opsin, Chrimson, is expressed in candidate IN1 sensory input neurons. **c-g**, Averaged normalised (ΔF/F_0_) GCaMP6s fluorescence in IN1s before and after optogenetic stimulation of candidate sensory neurons: Control group (**c**), sugar sensing neurons, Gr64f (**d**), bitter sensing neurons, Gr66a (**e**), water sensing neurons, ppk28 (**f**), and mechanosensory neurons, TMC (**g**). The optogenetic stimulation period is shown in a magenta-shaded area (mean ± SEM, stimulation = 1s, continuous, power = ∼0.75mW). (See also Extended Data Figs. 1a-g). **h**, Averaged normalised peak responses of IN1s in indicated genotypes upon optogenetic stimulation (n = 5-7 flies and five trials per fly, mean ± SEM, one-way ANOVA with Bonferroni post hoc test, \*\**p < 0.01*, not significant (n.s.)). IN1s responded to the activation of sugar-sensing neurons but not to other sensory neurons. Although TMC neurons appeared to activate IN1s weakly, this response did not reach statistical significance.

**Fig.2.**
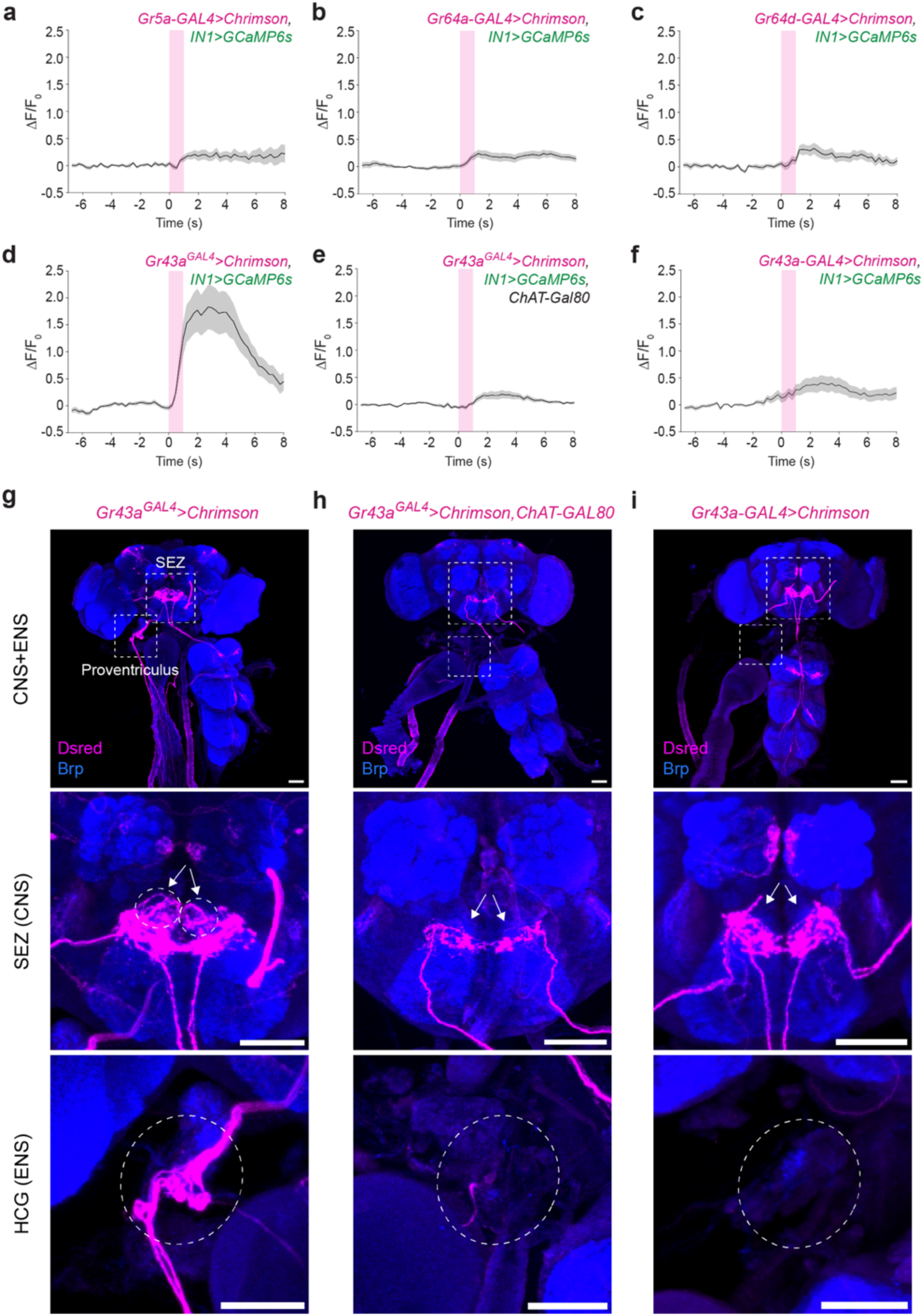
IN1s receive excitatory input from enteric sensory neurons expressing *Gr43a*. **a-c**, Averaged normalised (ΔF/F_0_) GCaMP6s fluorescence in IN1s before and after optogenetic stimulation of different classes of sugar-sensing neurons. IN1s do not respond to the activation of neurons expressing *Gr5a* (**a**), *Gr64a* (**b**), or *Gr64d* (**c**) (n = 5-7 male flies; mean ± SEM). The optogenetic stimulation period is shown in a magenta-shaded area (stimulation = 1s, continuous, power = ∼0.75mW out of objective). **d-f**, Stimulation of different classes of Gr43a neurons results in different responses in IN1s. IN1s are strongly activated only when enteric neurons express Chrimson (n = 6 male flies, mean ± SEM). (See also Extended Data Figs. 2a-d). **g-i**, Expression patterns of Gr43a transgenic flies expressing Chrimson (magenta) in the CNS+ENS (top), SEZ (middle) and HCG (ENS) (bottom). (**g**) *Gr43a^GAL^*^4^ knock-in labels enteric sensory neurons in the HCG and strongly activates IN1s (**d**). (**h**) Combining *Gr43a^GAL^*^4^ with *ChAT-Gal80* suppresses the expression in HCG and the responses in IN1s (**e**). (**i**) *Gr43a-GAL4* transgenic strain does not label enteric neurons and does not activate INs (**f**) (scale bars = 50 μm, white circles indicate enteric neuron cell bodies in HCG, white arrows indicate their projections in the SEZ.)

### Enteric Gr43a neurons penetrate the gut lumen and monitor sucrose ingestion

Since our results suggest that IN1s receive excitatory input from enteric Gr43a neurons, we hypothesised that these neurons would also respond to sucrose ingestion similarly to IN1s. Gr43a is one of the most conserved insect taste receptors specifically activated by the monosaccharide fructose^49^. It is expressed not only in chemosensory neurons but also present in other organs such as the brain and digestive tract^45^ (Fig. 2g). Although Gr43a is a fructose receptor^49^, neurons expressing Gr43a have been shown to respond to multiple sugars, including sucrose^43, 45^. To test our hypothesis, we captured the activity of enteric Gr43a neurons during sucrose ingestion. Since the cell bodies of these enteric neurons are located in the HCG next to the proventriculus^45^, we developed a novel preparation that allowed us to gain optical access to these neurons (Figs. 3a, b). By combining rapid volumetric two-photon imaging with our new surgical preparation, we successfully recorded the activity of enteric neurons *in vivo* during ingestion (Supplementary Video 1). In these experiments, some enteric Gr43a neurons (cell count=∼3-4) rapidly responded to the ingestion of high-concentration sucrose (∼1M), measured by GCaMP6s fluorescence in their cell bodies. In contrast, other enteric Gr43a neurons were tonically active throughout the imaging session (cell count=∼2-3) (Figs. 3c, d). The activity of sugar-responsive Gr43a neurons was transient, remaining elevated only during the period of ingestion (Fig. 3e). We then investigated whether Gr43a neural responses to sugar ingestion are dependent on the metabolic state or sugar concentration by recording their activity in fed and fasted flies offered high-concentration (∼1M) or low-concentration (∼100mM) sucrose. The sugar-evoked peak activity of Gr43a neurons was similar across all conditions (Figs. 3e, f). However, Gr43a neural activity persisted longer when flies consumed high-concentration sucrose compared to low-concentration (Figs. 3e, g). Interestingly, the persistent activity of Gr43a neurons is positively correlated with sugar concentration but is independent of the flies’ metabolic state. These results indicate that a subset of enteric Gr43a neurons monitor the sugar content of food during ingestion and convey this information to IN1s within seconds.

**Fig.3.**
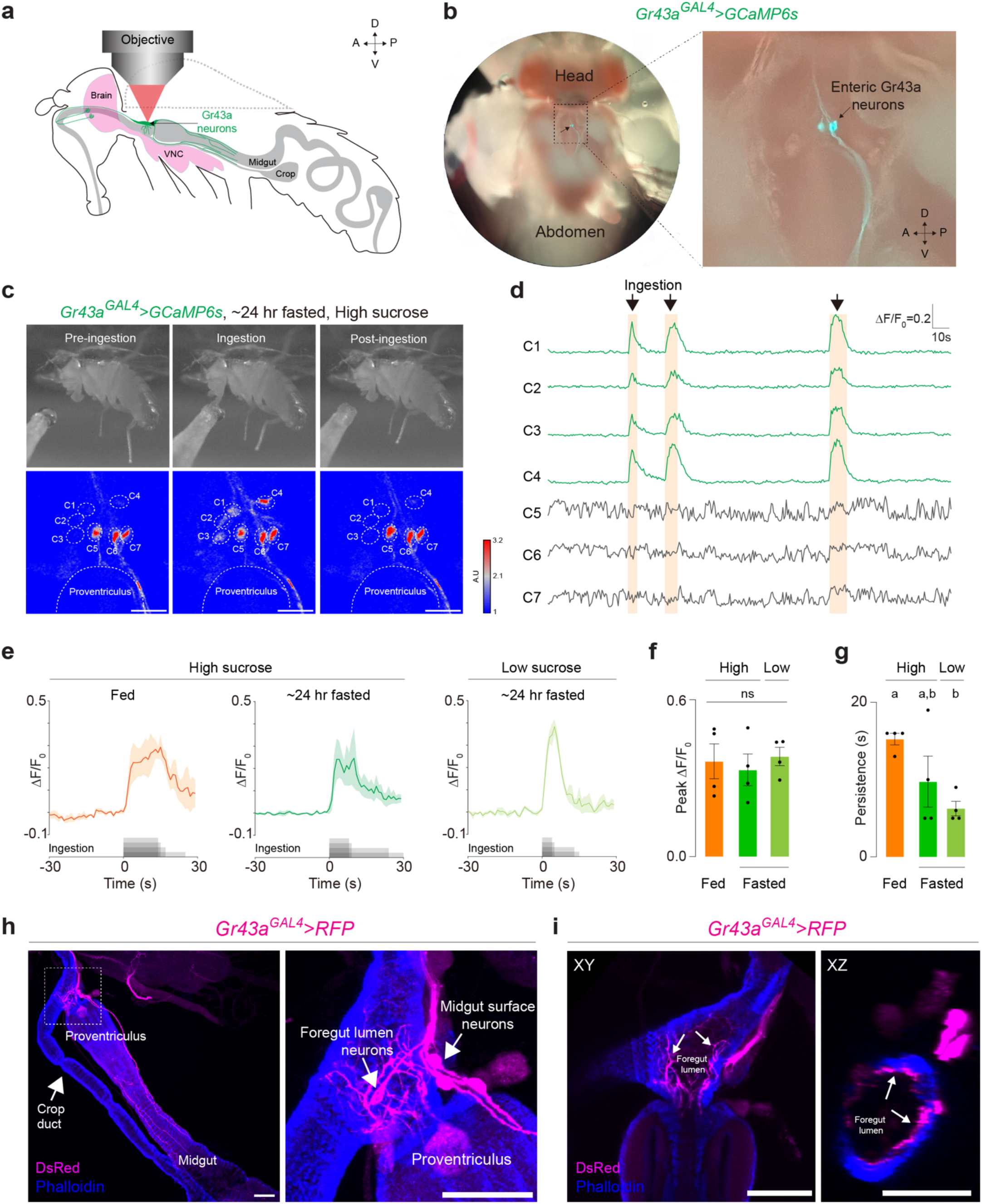
Enteric sensory neurons expressing *Gr43a* respond to sugar ingestion. **a** Schematic showing the enteric neuron imaging prep. Anterior (A), posterior (P), dorsal (D), ventral (V). The dashed line indicates the tissue removed. **b**, Top view of the enteric Gr43a sensory neurons expressing *GCaMP6s*. **c**, Representative GCaMP6s responses of Gr43a enteric sensory neurons recorded in the same 24-hr-fasted male fly before (left), during (middle), and after (right) ∼1M sucrose ingestion. Still images were captured by a video camera (top). Heatmap of Gr43a enteric sensory neuron activity in response to sucrose ingestion (bottom). Dashed white circles indicate seven separate cell bodies (C1-C7) of enteric Gr43a neurons (A.U., arbitrary units). **d**, Normalized (ΔF/F_0_) GCaMP6s fluorescence in individual Gr43a cell bodies. Neurons that respond to ∼1M sugar ingestion (C1-C4) are coloured in green; neurons that are not activated during sugar ingestion (C5-C7) are coloured in grey. The sugar ingestion period is shown as an orange-shaded area. **e**, Normalised (ΔF/F_0_) GCaMP6s fluorescence in active Gr43a enteric sensory neurons in response to high (∼1M and low sucrose (∼100mM) ingestion in fed and ∼24-hour fasted conditions (n = 4 male flies, mean ± SEM). The sugar ingestion is shown in grey. **f**, Averaged normalised (ΔF/F_0_) peak responses of Gr43a enteric sensory neurons in indicated conditions (n =4 male flies, mean ± SEM, one-way ANOVA, p = 0.8083). **g**, Persistence of Gr43a normalised (ΔF/F_0_) responses in indicated conditions (n =4 male flies, mean ± SEM, one-way ANOVA with Bonferroni post hoc test, p < 0.05). **h**, Detailed anatomical visualisation of enteric Gr43a neurons (magenta). A subset of Gr43a enteric sensory neurons penetrate the foregut and arborise in the inner surfaces of the gut lumen (foregut lumen neurons). Other Gr43a enteric sensory neurons do not penetrate the foregut and send their projections to the midgut (midgut surface neurons) (scale bars = 50μm) (See also Extended Data Figs. 3j-k). **i**, Cross-sectional images of Gr43a foregut lumen neurons (magenta) in different axes: XY (left) and XZ (right). Arrows indicate enteric Gr43a neurites penetrating the gut lumen (scale bars = 50μm).

Next, we examined the anatomical differences among enteric Gr43a neurons that might explain their distinct responses to sucrose ingestion. To do this, we acquired high-resolution images of Gr43a neural processes along the fly digestive tract (Figs. 3h, i, Extended Data Figs. 3j, k). Our morphological analysis revealed two classes of enteric Gr43a neurons: those that penetrate the foregut lumen at the junction of the crop duct and proventriculus (foregut lumen neurons) (Figs. 3h, i) and those that send projections along the midgut muscles (midgut surface neurons) (Extended Data Figs. 3j, k). The foregut lumen Gr43a neurons are ideally positioned to detect ingested sucrose, as their processes have direct access to the nutrients within the gut lumen (Figs. 3i). In contrast, midgut surface Gr43a neurons innervate the gut muscles, and their processes do not penetrate the midgut muscles (Extended Data Figs. 3k). Our findings demonstrate that foregut lumen Gr43a neurons directly monitor the sugar content of food within the gut lumen and relay this information to IN1s. This rapid sensory feedback mechanism enables flies to assess the nutrient content of food as they ingest it, allowing them to decide whether to continue or stop eating.

### IN1s receive excitatory input from two classes of enteric sensory neurons expressing *Gr43a*

Based on our anatomical and functional analysis, we propose that Gr43a neurons that penetrate the foregut lumen respond to sucrose ingestion and activate IN1s in the brain. To investigate this further, we generated split-GAL4^50, 51^ lines targeting distinct classes of Gr43a neurons using enhancers expressed in the enteric nervous system (Extended Data Figs. 3a-i). Split-GAL4s were first screened for expression in the enteric nervous system, then used in functional connectivity analysis using optogenetics coupled with two-photon functional imaging (Fig. 4a). Out of the 20 split-GAL4 lines generated, 15 labelled enteric sensory neurons, and, of these 7, significantly activated IN1s upon optogenetic stimulation (Fig. 4b, Extended Data Figs. 4a-o). Next, we explored whether all enteric neurons capable of activating IN1s also respond to sucrose ingestion. Only enteric neurons labelled by *EN-13>* were activated by sucrose ingestion. In contrast, the others showed no sugar-evoked responses (Fig. 4d). Our anatomical analysis revealed that *EN-13>* labels enteric Gr43a neurons whose processes penetrate the gastrointestinal tract (foregut lumen neurons) enabling them to detect sucrose within the gut lumen during ingestion (Figs. 4e, f). These findings support our hypothesis that Gr43a neurons in the foregut lumen can monitor the sucrose content of ingested food and send positive feedback signals to IN1s to sustain sugar intake. Interestingly, we have identified other enteric sensory neurons that can induce IN1 activity, yet they do not respond to sugar ingestion. These neurons may detect other nutrients or mechanical stimuli in the gut lumen, potentially regulating various aspects of feeding behaviour or metabolic processes.

**Fig.4.**
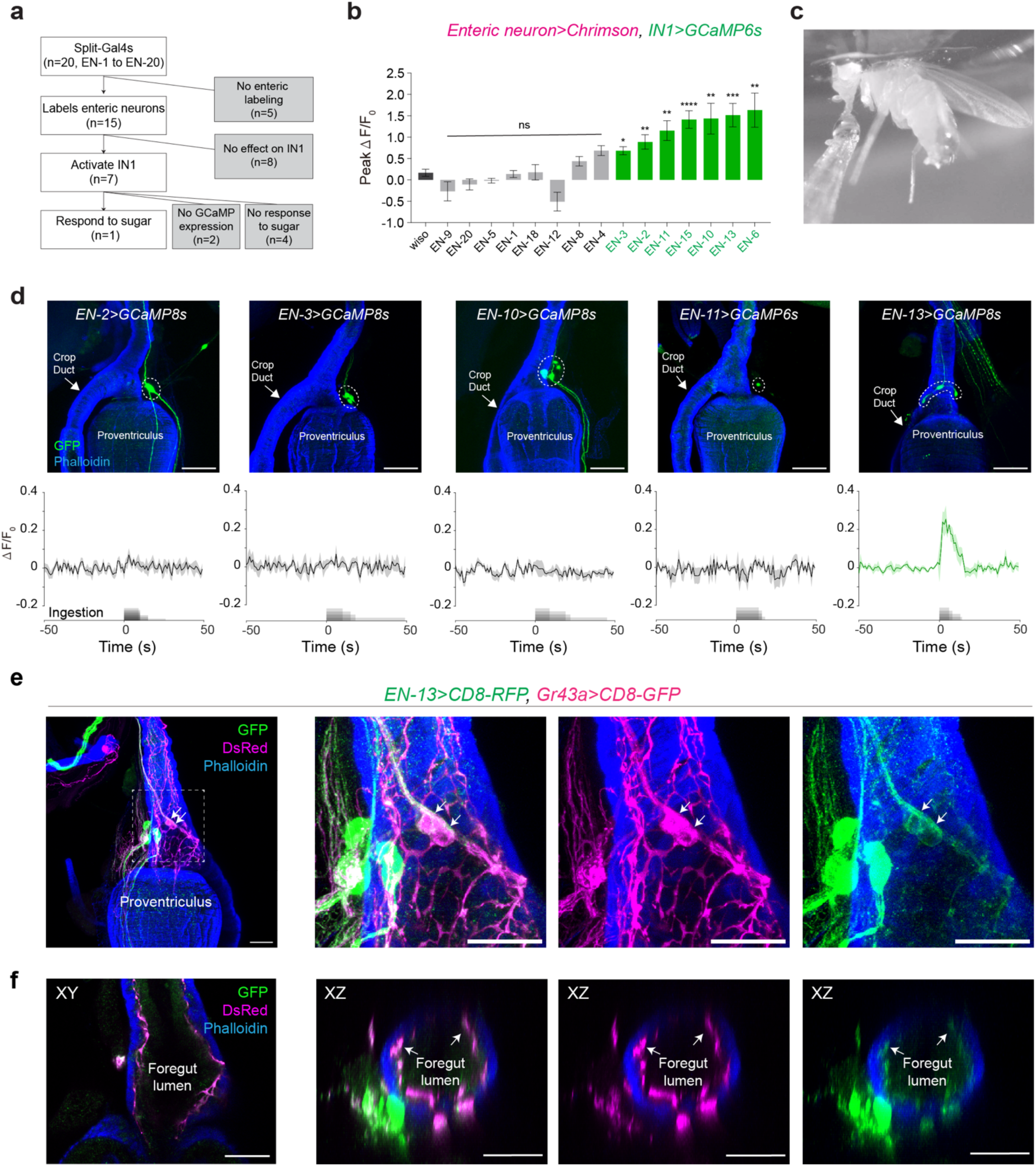
Different classes of enteric sensory neurons can activate IN1s. **a**, Overview of the enteric sensory neuron two-photon functional imaging screen. We generated 20 split-GAL4s (see also Extended Data Figs. 3a-i). Fifteen of these split-GAL4s labelled different populations of enteric sensory neurons. Optogenetic activation of seven enteric split-GAL4s (*ENs>*) activated IN1s, and one of these lines also responded to sugar ingestion. **b**, Averaged normalised (ΔF/F_0_) peak responses of IN1s upon optogenetic stimulation of enteric split-GAL4s (*ENs>*). Positive ENs are shown in green; negative ENs are shown in grey; the control group is shown in black. (n = 5-7 male flies and five trials per fly, mean ± SEM, Kruskal Wallis test with Dunn’s post hoc test, \**p < 0.05*, \*\**p < 0.01*, \*\*\**p < 0.001*, \*\*\*\**p < 0.0001*) (see also Extended Data Fig. 4). **c**, A representative image showing enteric sensory neuron imaging during sugar ingestion. A male fly is fixed from its thorax underneath the imaging objective. Sugar stimulus is delivered using a pulled glass pipette. **d**, Confocal images showing the expression of GCaMP6s or GCaMP8s in each *ENs>* (green). Dashed white circles indicate the ROIs used to quantify the GCaMP6s fluorescence (scale bars = 50μm.) (top). Normalised (ΔF/F_0_) GCaMP6s fluorescence in *ENs>* in response to high sucrose (∼1M) ingestion in 24-hour fasted flies (n = 4-7 male flies; mean ± SEM), with sugar ingestion shown in grey (bottom). **e**, Immunohistochemical analysis of neurons labelled by *EN-13>* (magenta) and *Gr43a*> (green) in the HCG. White arrows indicate the enteric sensory neurons labelled by both transgenic lines (scale bars = 25μm). **f**, Cross-sectional images of neurons labelled by *EN-13>* (magenta) and *Gr43a*> (green) in different axes: XY (left) and XZ (middle-right). Arrows indicate enteric sensory arbours penetrating the gut lumen (scale bars = 25μm).

### IN1s receive synaptic input from enteric afferents and are anatomically distant from feeding initiation circuits

Next, we used connectomics to characterise the anatomical organisation of IN1s and their interactions with different classes of gustatory receptor neurons (GRNs) and neurons that regulate feeding initiation. Recently, the whole-brain connectome of an adult fly has been completed and released to the *Drosophila* community through the online FlyWire platform, describing the synaptic organisation of ∼130,000 neurons^52–55^. Flywire uses the full adult female brain (FAFB) dataset, the first electron microscopy (EM) volume of an adult fly brain^56^, which contains the neurons in the primary taste processing centre of the fly brain, SEZ^57, 58^. FAFB volume has recently been segmented automatically, allowing computer-based detection of synapses^54, 59^. Using the FAFB connectome, we first identified four putative IN1s (IN1-1, IN1-2, IN1-3, and IN1-4) based on their cell body locations, projection patterns, and synaptic organisations (Extended Data Figs. 5a, b). Synaptic network analysis revealed that putative IN1s have a similar number of presynaptic (n=825±20) and postsynaptic (n=585±32) connections. Interestingly, we found that IN1s are recurrently connected to each other (Extended Data Fig. 5d). This synaptic organisation might indicate feed-forward excitation among IN1s, which could play a crucial role in generating their persistent activity upon sugar ingestion.

Using the connectome, we further investigated the synaptic distance of IN1s to different classes of GRNs that are annotated in the FlyWire data analysis platform Codex (Connectome Data Explorer: codex.flywire.ai)^58, 60^ (Extended Data Fig. 5e). Our analysis demonstrated that IN1s do not receive direct synaptic input from any of the labellar GRNs or the majority of pharyngeal GRNs. (Extended Data Figs 5g-i). Since our functional connectivity experiments revealed that IN1s receive functional input from enteric Gr43a neurons (Figs. 2, 3), we next investigated whether enteric afferents provide direct synaptic input to IN1s. We first annotated putative enteric afferents in the FAFB connectome based on their characteristic projections in the prow area of the SEZ (Extended Data Figs. 5e, f). We found that several putative enteric afferents provide direct synaptic input to IN1s (Extended Data Fig. 5f), further supporting their role in regulating food ingestion rather than initiating feeding behaviour.

We next investigated the connectivity between IN1s and neurons that regulate feeding initiation. Recent studies have identified neural circuits in the adult fly brain governing this behaviour. These sensorimotor circuits connect GRNs to motor neurons that innervate the proboscis muscles^57, 61^. Most neurons in this pathway respond to sugar ingestion and can induce proboscis extension, a behaviour associated with feeding initiation^57, 61^. Our analysis showed no direct synaptic connections between IN1s and second-order, third-order or pre-motor neurons that regulate feeding initiation (Extended Data Figs. 6a, b). Overall, our connectome analysis revealed that IN1s are synaptically distant from sensory and central neurons regulating feeding initiation. Instead, as our functional connectivity analysis also demonstrated (Figure 4), they receive direct synaptic input from enteric sensory neurons linked to food ingestion.

### IN1s are synaptically connected to local SEZ neurons

Since IN1s are not directly connected to neurons that regulate feeding initiation, we asked which circuits they are connected to in the fly brain. All four putative IN1s (IN1-1, IN1-2, IN1-3, and IN1-4) received presynaptic and postsynaptic connections from a comparable number of neurons (n=55.75±1.4 and n=29.75±1.1 respectively). Interestingly, most of these neurons were intrinsic to SEZ, with few exceptions that project outside this region (Extended Data Figs. 7a-h, Extended Data Figs. 8a-h). We first characterise the IN1 presynaptic neurons. Our connectomics analysis showed that four local SEZ neurons contributed the highest count of input synapses to IN1s. PRW.10 contributed the most, with 82.5±4 synapses per IN1, followed by PRW.70, with an average of 63.75±3, PRW.9, with an average of 49.25±1.6, and PRW.GNG.8 with an average of 47.75±5 synapses (Extended Data Figs. 7b, d, f, h). All of these IN1 presynaptic neurons are predicted to be cholinergic^62^, and they have processes within the prow region of the SEZ (Extended Data Figs. 7i). Next, we examined IN1 postsynaptic neurons. Similar to presynaptic inputs, the primary outputs of IN1s appear to be neurons with cell bodies within the SEZ: PRW.248 (54±13 synapses), PRW.249 (63±4 synapses), PRW.263 (82.5±2.5), PRW.274 (55.5±13.5), PRW.281 (67.5±2.5), PRW.310 (57.5±2.5) and Doublescoop^63^ (55±4) (Extended Data Figs. 8a-h). PRW.248, PRW.249, PRW.263, PRW.274, PRW.281, and PRW.310 belong to the same class of SEZ neurons (previously dubbed “Peep” neurons^63^) with dendrites located in the prow and no evident axons within the brain, suggesting they are putative motor neurons. These neurons represent the primary synaptic output of IN1s, accounting for 28-35% of their output synapses. The second class of IN1 output neurons consist of those previously dubbed as Doublescoop neurons^63^. Doublescoop neurons have cell bodies in mediolateral SEZ and send projections towards the midline, where IN1 processes are located (Extended Data Figs. 8b, d, f & h). These neurons are predicted to be cholinergic^62^, and like the IN1 presynaptic neurons, their processes are intrinsic to the SEZ. Our connectome analysis revealed that IN1s’ main synaptic inputs and outputs are local SEZ neurons, most of which have not been previously described. Furthermore, the major synaptic output of IN1s are motor neurons that project outside of the brain (Extended Data Fig. 8i). We hypothesised that IN1 motor output neurons are most critical for their functions in regulating fly ingestion, leading us to focus our further investigation on these neurons.

### IN1s inhibit the activity of enteric motor neurons that innervate the crop duct

To further investigate the relationship between IN1s and their motor output neurons (PRW.248, PRW.249, PRW.263, PRW.274, PRW.281, and PRW.310), we first generated split-GAL4^50, 51^ lines to gain genetic access to them (Figs. 5a, b). Our anatomical characterisation verified that these neurons extend long projections outside the brain to the gastrointestinal tract, where they innervate the entry segments of the crop duct. (Fig. 5c). We renamed these neurons as Crop-innervating Enteric Motor (CEM) neurons to more precisely reflect their anatomical organisation. CEM neurons do not appear sexually dimorphic and exist in both male (Fig. 5b) and female brains (Extended Data Figs. 9a, b). The axons of these neurons project outside the brain along the oesophagus and branch at the junction between the crop duct and the entry of the proventriculus (Figs. 5b, c, Extended Data Figs. 9a, b). Immunohistochemical analysis showed that the synaptic boutons of CEM neurons express the vesicular glutamate transporter (VGLUT), confirming they are indeed glutamatergic motor neurons (Fig. 5c). To further examine the functional connectivity between CEM neurons and IN1s, we performed two-photon functional imaging coupled with optogenetic stimulation. Optogenetic activation of IN1s inhibited the activity of CEM neurons. This inhibitory effect was modest during the short (1-second) optogenetic trials but became more pronounced in longer (10-second) ones (Figs. 5d, e). Our results demonstrated that IN1s provide inhibitory synaptic input to CEM neurons.

**Fig.5.**
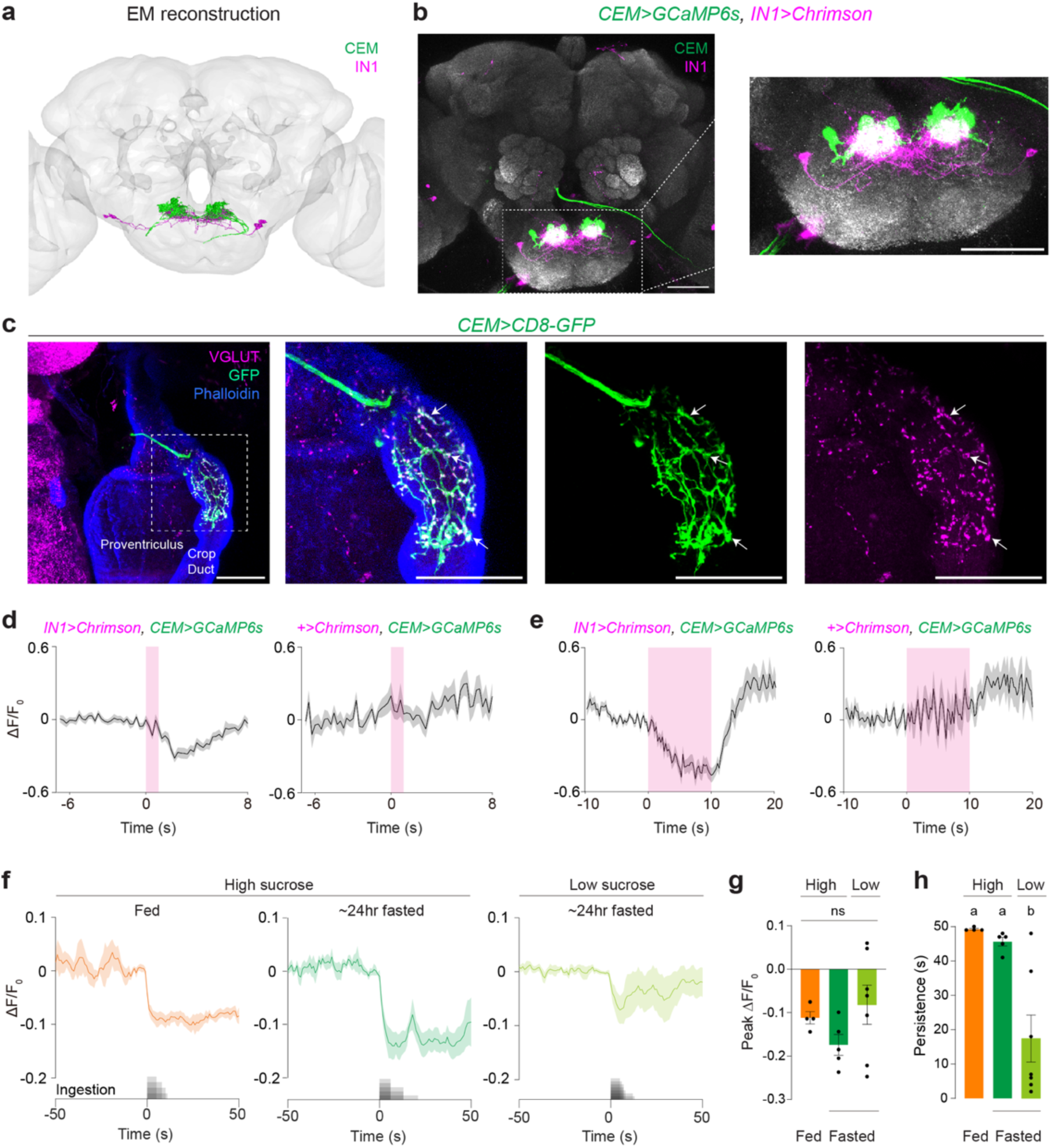
IN1s inhibit CEM neurons upon sucrose ingestion. **a**, Electron microscopy (EM) reconstruction of putative INs (magenta) and the CEM neurons (green). **b**, Confocal images of IN1s (magenta) and CEM neurons (green) in the brain (left). IN1 and CEM arbours are intermingled in the SEZ (right) (scale bars = 50μm). **c**, Staining of CEM axonal terminals with VGLUT and GFP antibodies in *CEM>CD8-GFP* flies. White arrows indicate the co-labelling of CEM synaptic terminals in the crop duct with VGLUT (magenta) and GFP (green) (scale bars = 25μm). **d-e**, Optogenetic stimulation of INs inhibits the activity of CEM neurons. (n = 5-8 male flies; mean ± SEM). The optogenetic stimulation period is shown in a magenta-shaded area (stimulation = 1s (**d**) or 10s (**e**), continuous, power = ∼0.75mW). **f**, Normalised (ΔF/F_0_) GCaMP6s fluorescence in CEM neurons in response to high (∼1M) and low (∼100mM) sucrose ingestion in fed and ∼24-hour fasted conditions (n = 4-7 male flies; mean ± SEM). The sugar ingestion is shown in grey. **g**, Averaged normalised (ΔF/F_0_) peak responses of CEM neurons in indicated conditions (n = 4-7 male flies, mean ± SEM, one-way ANOVA, p = 0.2353). **h**, Persistence of CEM normalised (ΔF/F_0_) responses in indicated conditions (n =4-7 male flies, mean ± SEM, one-way ANOVA with Bonferroni post hoc test, p < 0.05).

Next, we investigated the activity of CEM neurons during sugar ingestion. Given that IN1s are excited by sugar ingestion and exhibit state- and stimulus-specific responses^8^, we hypothesised that CEM neurons might reflect IN1 activity and would be inhibited by sugar ingestion due to their inhibitory synaptic inputs from IN1s. Supporting this hypothesis, our functional imaging experiments showed that CEM neurons were persistently inhibited upon the ingestion of high-concentration sucrose in both fed and 24-hour-fasted conditions (Fig. 5f, Supplementary Video 2). The inhibitory response to sucrose was transient when 24-hour-fasted flies consumed low-concentration sucrose (Fig. 5f). Our quantitative analysis revealed that the peak response in CEM neurons was similar across all conditions (Fig. 5g). However, we observed that the persistence of inhibition was significantly reduced in 24-hour-fasted flies consuming low-concentration sucrose compared to those consuming high-concentrations (Fig. 5h). Our findings revealed an inverse correlation between the activity of CEM neurons and IN1s: IN1s remain persistently activated during the ingestion of high-concentration sucrose^8^, while CEM neurons are continuously inhibited. Moreover, this persistent activation of IN1s^8^ and inhibition of CEM neurons depend on sugar concentration, occurring only when flies ingest high-concentration sucrose but not low-concentration.

### IN1 ingestion circuit mediates sugar intake by controlling food entry to the crop

In *Drosophila* and other insects, ingestion is regulated by a series of peristaltic muscle contractions that pump the food into the gastrointestinal tract^3, 64, 65^. After the ingested food passes through the oesophagus, it reaches the crop duct and proventriculus, where it must be sorted to either enter the crop, a stomach-like storage organ or proceed to the midgut through the proventriculus. Mosquitoes transport meals with low sugar content directly to the midgut, whereas sugar-enriched meals are stored in the crop^66, 67^. However, the neural circuits that regulate sugar transport within the gastrointestinal tract are unknown. We hypothesised that the IN1 ingestion circuit might regulate sugar intake by mediating the transport of sugar-enriched foods into the crop. To test this hypothesis, we first investigated whether flies regulate the transport of ingested food based on its sugar content, similar to other insects^68,69,70^. We developed an *in vivo* imaging assay to monitor the food entry into different gastrointestinal compartments in body-fixed flies. In this assay, a fluorescent dye, fluorescein, was mixed with high and low-concentration sucrose and fed to 24-hour fasted flies. Using *in vivo* two-photon imaging, we then tracked the movement of the fluorescent food within the oesophagus and crop during and after ingestion. (Fig. 6a, Extended Data Fig. 10a). When flies ingested high-concentration sucrose, the crop duct remained persistently open, as indicated by the sustained fluorescent signal, allowing continuous food passage into the crop. In contrast, when they consumed low-concentration sucrose, the crop duct opened only briefly, resulting in a short-lived fluorescent signal (Figs. 6b, c). These differences in food transport were not apparent in the oesophagus (Extended Data Figs. 10b, c), indicating flies can modulate the transport of food to their crop based on its sugar content.

**Fig.6.**
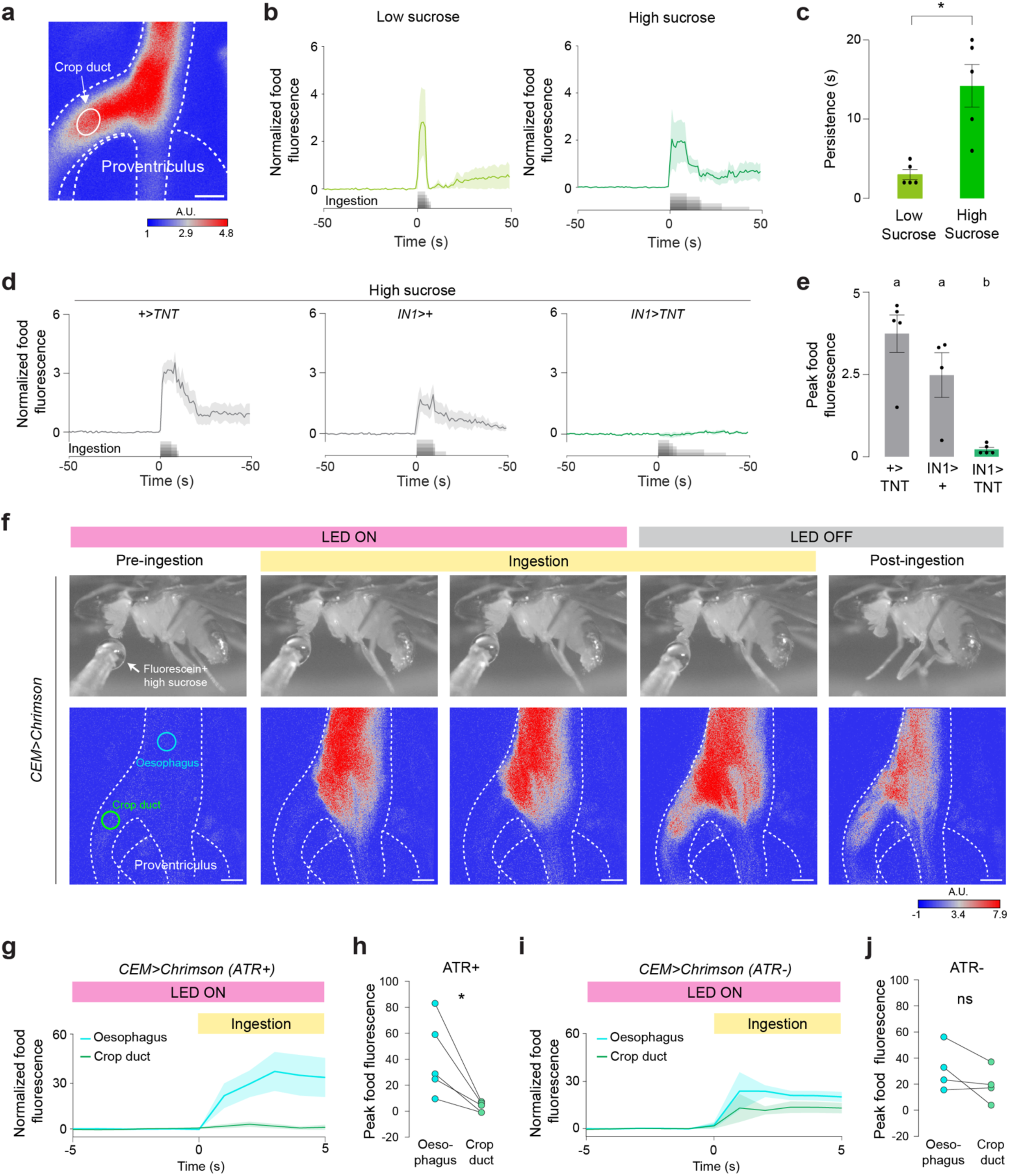
CEM neurons gate the entry of sucrose into the crop during ingestion. **a,** A representative image showing the ROI in the crop-duct during ingestion of high-sucrose containing fluorescein (scale bar = 25μm. A.U., arbitrary unit). **b**, Normalized food fluorescence in the crop duct before and after ingestion of low (∼100mM) (left) and high (∼1M) (right) sucrose (n = 5 male flies, mean ± SEM). The sugar stimulus was provided *ad libitum*. **c**, Persistence of normalised food fluorescence when flies are ingesting low (∼100mM) or high (∼1M) sucrose (n = 5 male flies, mean ± SEM, Unpaired t-test with Welch’s correction, \**p < 0.05*). **d**, Normalized food fluorescence in the crop duct before and after ingestion of high sucrose (∼1M) in flies with indicated genotypes (n = 4-5 male flies, mean ± SEM). **e**, Peak food fluorescence in the crop duct after ingesting high sucrose (∼1M) in flies with indicated genotypes. The food volume that enters the crop duct is significantly reduced when IN1 neurons are inhibited (n = 4-5 male flies, mean ± SEM, one-way ANOVA with Bonferroni post hoc test, p < 0.05). **f**, Representative images (top) and food fluorescence in the crop duct and oesophagus (bottom) of a 24-hr-fasted *CEM>Chrimson* male fly before (pre-ingestion), during (ingestion), and after (post-ingestion) high sucrose ingestion, and with and without optogenetic stimulation of CEM neurons. High sucrose (∼1M) solution cannot enter the crop duct until the CEM optogenetic activation is turned OFF (LED OFF). Cyan and green circles indicate ROIs in the oesophagus and crop duct, respectively (scale bars = 25μm, A.U., arbitrary unit). **g-j,** Normalized food fluorescence in the oesophagus (cyan) and crop duct (green) of *CEM>Chrimson* flies during optogenetic stimulation with ATR (**g**, test group) and without ATR (**i,** control group). (**h**) Optogenetic stimulation reduces the amount of sucrose that can enter the crop duct in the test group quantified by peak food fluorescence. (**j**) However, control flies are not affected by optogenetic stimulation (n = 4-5 male flies, mean ± SEM, paired t-test, ns, \**p<0.05*).

Building on our findings, we then tested whether the activity of IN1s or CEM neurons is required for sugar transport into the crop. Using two-photon imaging, we monitored the flow of high sucrose and fluorescein mixture to different digestive compartments while manipulating the activity of these neurons. Inhibiting synaptic vesicle release by expressing tetanus toxin light chain^68^ in IN1s (*IN1>TNT*) blocked the entry of high-concentration sucrose into the crop (Figs. 6d, e) without affecting the food transport to the oesophagus (Extended Data Figs. 10d, e). These results explain why *IN1>TNT* flies can only consume small volumes of food, as we previously demonstrated^8^. We then manipulated the activity of CEM neurons to determine if their activity is also required for sugar transport to the crop. Since IN1s inhibit CEM neurons during sugar ingestion (Fig. 5f), we hypothesised that activating CEM neurons during ingestion might mimic the effects of IN1 inhibition. To test this, we expressed the red-shifted channelrhodopsin Chrimson^25, 28^ in CEM neurons and photoactivated them during ingestion. Under continuous optogenetic stimulation, flies were unable to transport sucrose into their crop (Figs. 6g, h, Supplementary Video 3). However, once the optogenetic stimulation stopped, sucrose was able to enter the crop, allowing the flies to resume ingestion (Supplementary Video 3). Importantly, this effect on food ingestion was not due to red light stimulation, as control flies of the same genotype that were not fed all-trans-retinal (ATR), a co-factor essential for Chrimson activity, showed no impairments in food transportation from the oesophagus to the crop (Figs. 6i, j, Supplementary Video 3). Our findings indicate that the coordinated activity of IN1s and CEM neurons is essential for transporting sugar-rich foods into the crop and thereby optimising flies’ ingestive behaviours. When the activity of these neurons is disrupted, ingested food cannot be moved into the crop. This impairment restricts flies’ ability to consume and store large volumes of sugar, highlighting the critical role of this gut-brain-gut interoceptive circuit in controlling sugar ingestion.

## Discussion

Gut-brain circuits have been linked to nutrient preference and food ingestion in humans^69, 70^, rodents^17, 18, 20, 21, 24^, and insects^5, 9–13, 65^. In mammals, these circuits sense nutrients^16, 18, 21^ or stretch^14, 24^ within the stomach or intestinal lumen and send feedback signals to central and motor circuits, which mediate swallowing, digestion and gut motility^2, 6, 22–24^. Here, we reveal a gut-brain-gut interoceptive circuit that regulates state and concentration-specific sugar ingestion in *Drosophila*. Our data support a model in which sugar-responsive enteric sensory neurons in the gut provide real-time nutrient information to the brain, specifically to IN1s that are synaptically connected to enteric motor neurons. When food-deprived flies consume sugar-rich foods, these enteric sensory neurons rapidly convey the sugar stimulus to IN1s, leading to their persistent activation and, consequently, inhibition of enteric motor neurons that innervate the crop duct muscles. This coordinated activity allows flies to open their crop duct and continuously transport food from the oesophagus into the crop, thereby enhancing their capacity to ingest and store large volumes of sugar-rich foods. The dynamic regulation of this interoceptive circuit by metabolic state and sugar concentration is crucial for flies to quickly assess the nutritional value of ingested foods and adjust their digestive processes accordingly, either stimulating or halting their food intake as needed. We propose that this gut-brain-gut interoceptive circuit plays a crucial role in enabling flies to optimise their dietary intake by prioritising the ingestion and storage of foods enriched in sugar, which provides a quick and efficient energy source. This adaptive feeding behaviour is likely to contribute to the survival and fitness of flies by maximising their energy acquisition in environments where food quality fluctuates.

Our study reveals that the interoceptive gut-brain circuit we have identified here in flies closely parallels the vagal sensorimotor circuits that mediate gut-brain communication in mammals. In mice, vagal sensory neurons reside in the nodose ganglia, where they transmit nutrient-derived signals (e.g., sugars, fats) from the gut to the hindbrain, specifically targeting the brainstem nuclei, the nucleus of the solitary tract (NTS)^2, 6, 24^. Recently, vagal sensory neurons that stimulate sugar preference and intake have been identified in mice^17, 18^. In contrast, the cell bodies of vagal motor neurons are located in the dorsal motor nucleus of the vagus (DMV) in the brainstem. These neurons project to various regions of the gastrointestinal tract, including the stomach, small intestine, large intestine, gallbladder, and pancreas, where they exert control over digestive processes^71^. Our results demonstrate that in *Drosophila*, enteric sensory neurons located in the hypocerebral ganglion (HCG, analogous structure to mammalian nodose ganglia) detect sugars in the gut lumen and relay this information to the IN1s in the brain. INs are located in the subesophageal zone (SEZ) of the fly brain, which processes gustatory sensory information similarly to the mammalian brainstem, particularly the NTS. The major synaptic output of IN1s is enteric motor neurons that project to the digestive tract, innervating the entry segments of the fly crop, a stomach-like organ. Activation of this gut-brain-gut circuit opens crop muscles, allowing flies to ingest large volumes of sugar-rich foods. Our findings reveal striking anatomical and functional parallels between the vagal sensory and motor neurons in mice and the enteric sensory and motor neurons in flies, suggesting conserved neural mechanisms for processing gut-derived sensory signals across evolutionarily distinct species.

Another notable finding in our study is the demonstration that the IN1 ingestion circuit is anatomically distinct and synaptically distant from the feeding initiation circuits within the fly brain^57, 61^. Recent whole-brain imaging studies in flies have provided compelling evidence for the presence of a functional map within the SEZ^72^. Parallel investigations in mammalian models, particularly in mice, have similarly identified topographic and functional representations within the brainstem nuclei, NTS^73, 74^ and DMV^71^. Our connectome analysis of the IN1 gut-brain interoceptive circuit further supports the idea of functional segregation within feeding-related neural circuits in the fly brain. This separation suggests a hierarchical organisation, where distinct but interconnected neural circuits process each step of food intake, from sensory detection of food to feeding initiation and sustained ingestion. Understanding this interconnected network could reveal fundamental principles underlying feeding behaviour across species and provide insights into the conserved neural computations that regulate food intake and interoception in the brain. In the long term, this knowledge may pave the way for novel therapeutic strategies to treat human disorders related to gut-brain dysregulation, such as obesity and eating disorders.

## Author Contributions

X.C. and N.Y. designed the study. X.C. performed all experiments except for enteric neuron immunohistochemistry in Extended Fig. S3. M.R.M. identified and characterised the enteric neuron GAL4s from the Janelia FlyLight first-generation GAL4 collection. N.Y. and S.N.T. cloned the IN1 split GAL4 and split LexA plasmids and generated the transgenic lines. X.C. performed the analysis in Figs. 1-6 and Extended Data Fig. 2. X.C. performed the connectivity network analysis in Extended Data Figs. 5f-i and Extended Data Fig. 6b. X.C. annotated the IN1s in the Flywire connectome. N.Y. analysed the Codex data and generated the EM reconstructions in Extended Data Figs. 5-7. X.C. and N.Y. wrote the manuscript with input from M.R.M. and S.N.T.

## Acknowledgements

We thank Marco Gallio, Azahara Oliva, Gabriella Sterne, Yuhan Wang, and the members of the Yapici Lab for their comments on the manuscript. We thank Barry Dickson for sharing the ZpLexADBD-pBGUw pre-publication. We thank Tianxing Jiang for building the electronics to synchronise the video and functional imaging data acquisition and Haein Kim for helping us with immunohistochemistry experiments. We acknowledge Bloomington Drosophila Stock Center (NIH P40OD018537) for reagents. We extend our special thanks to the entire Flywire community for their invaluable contributions to proofreading the FAFB connectome. Especially to Greg Jefferis, who has helped us identify the IN1s in the FAFB connectome. This project was supported by the Cornell University Nancy and Peter Meinig Family Investigator Program, the Pew Biomedical Scholar Award, the Alfred P. Sloan Foundation Award, and the National Institutes of Health grant NIH-R35 (5R35GM133698) and NIH-R21 (R21AI149772).

## Data and code availability

The raw data and resource information that support the findings of this study are available from the corresponding author upon reasonable request. Source data are provided in this paper.

## Code availability

All analysis code will be available at https://github.com/Nilayyapici/Cui_et_al

## Reporting summary

Further information on research design is available in the Research Reporting Summary linked to this paper.

## Extended data

**Extended Data Fig.1.**
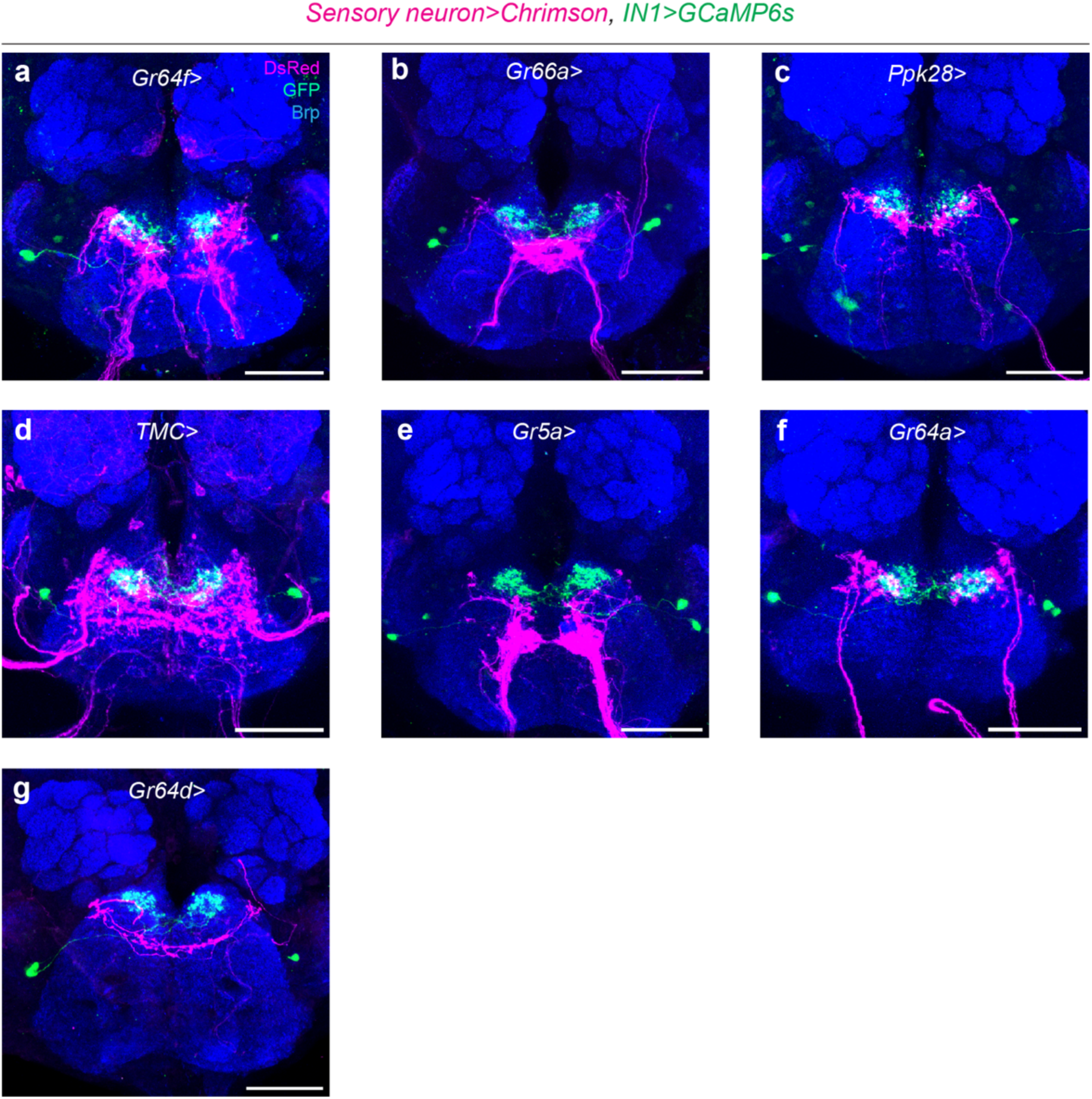
Expression patterns of GAL4 lines in SEZ labelling different classes of sensory neurons. **a-g**, Confocal images of sensory neuron afferents (magenta) and IN1 arbours (green) in the anterior SEZ. The sensory neurons are labelled by *Gr64f>* (**a**), *Gr66a>* (**b**), *ppk28>* (**c**), *TMC>* (**d**), *Gr5a>* (**e**), *Gr64a>* (**f**), *Gr64d>* (**g**). (scale bars = 50μm).

**Extended Data Fig.2.**
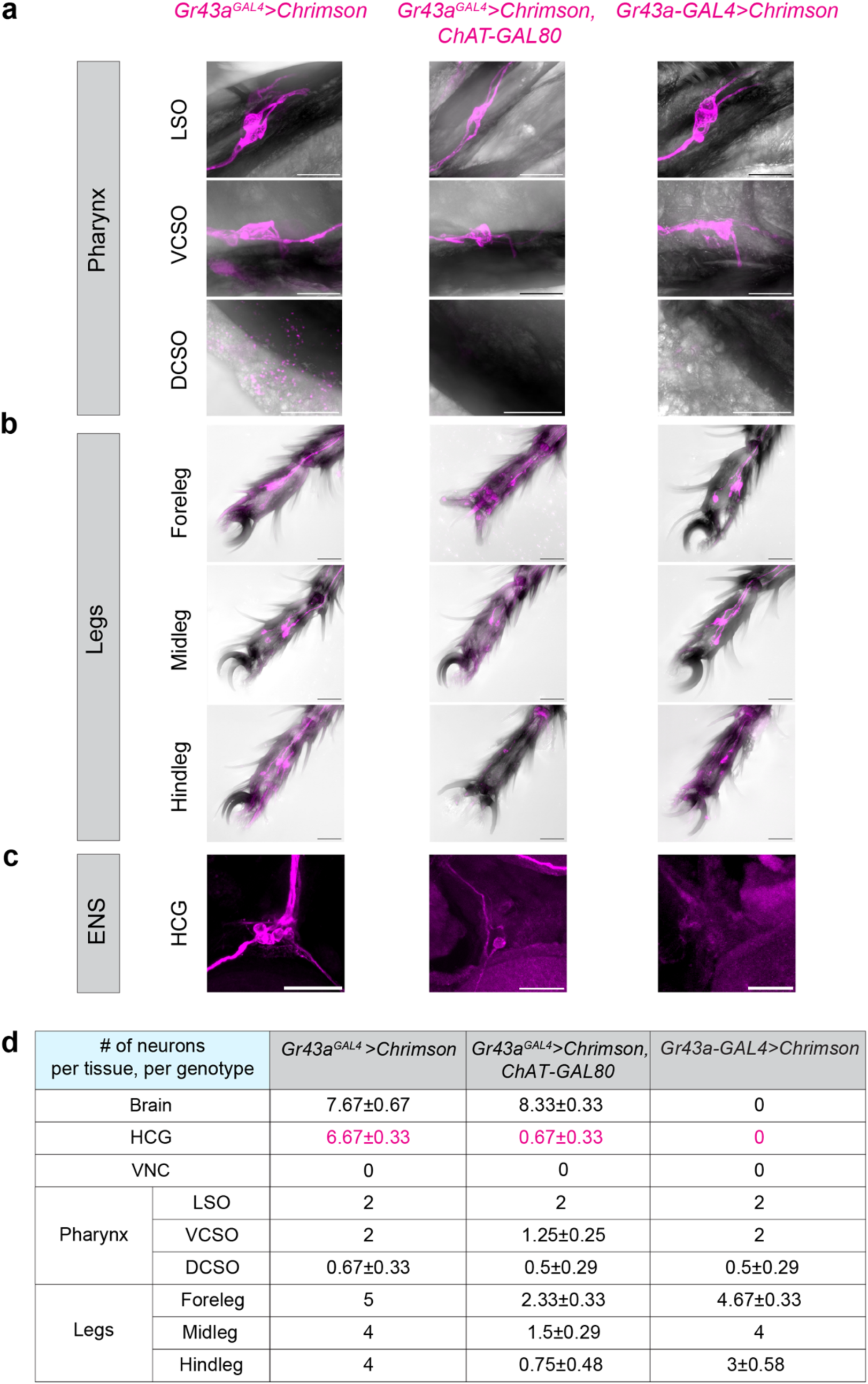
Expression patterns of Gr43a transgenic lines in the chemosensory and enteric neurons. **a-c**, Expression patterns of different transgenes labelling distinct classes of Gr43a neurons (magenta) in various chemosensory organs and the enteric nervous system (scale bars = 25μm). **d**, Quantification of Gr43a neurons in flies carrying the indicated transgenes in the CNS, ENS and various chemosensory organs (HCG, hypocerebral ganglion; VNC, ventral nerve cord; LSO, labral sense organ, VCSO, ventral cibarial sense organs, DCSO, dorsal cibarial sense organs). For legs and pharyngeal organs, LSO, VCSO and DCSO, we report the number of neurons unilaterally (n = 3-4 male flies per group, mean ± SEM).

**Extended Data Fig.3.**
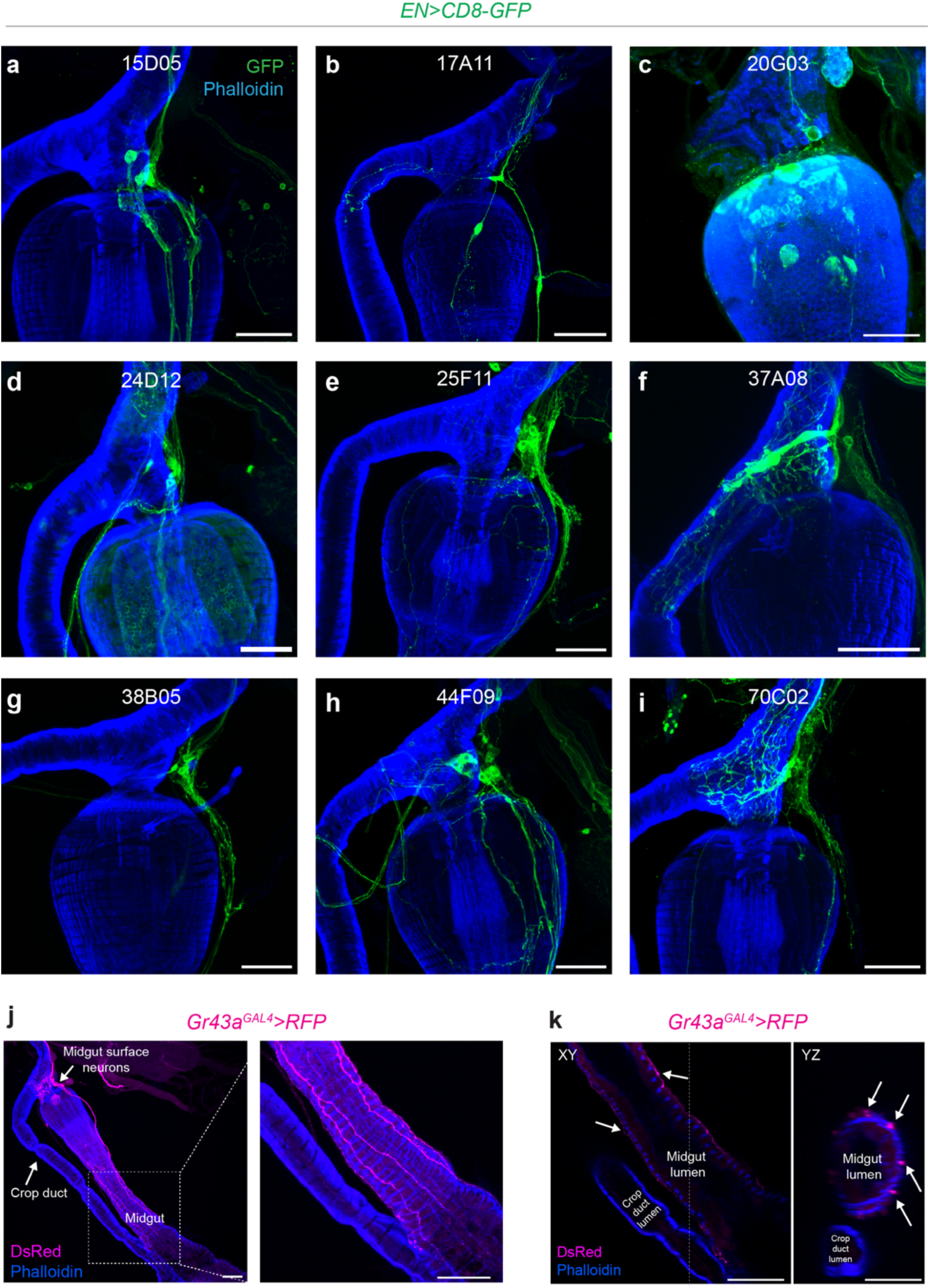
GAL4 lines labelling different classes of enteric neurons. **a-i**, Confocal images of enteric neurons (green) labelled by selected GAL4 lines whose promoters are used to generate the EN-split GAL4s (scale bars = 50μm). **j**, Confocal images of enteric Gr43a neurons that project to the midgut (magenta). The upper white arrow indicates the midgut surface neuron cell bodies, and the lower white arrow indicates the crop duct (left). Notice the midgut surface neurons arborise along the surface of the midgut muscle (blue) (right) (scale bars = 50μm) **k**, Cross-sectional images of Gr43a midgut surface neurons in different axes: XY (left) and XZ (right). Arrows indicate the neurites of Gr43a midgut surface neurons (magenta) innervating the gut muscles (blue) (scale bars = 50μm).

**Extended Data Fig.4.**
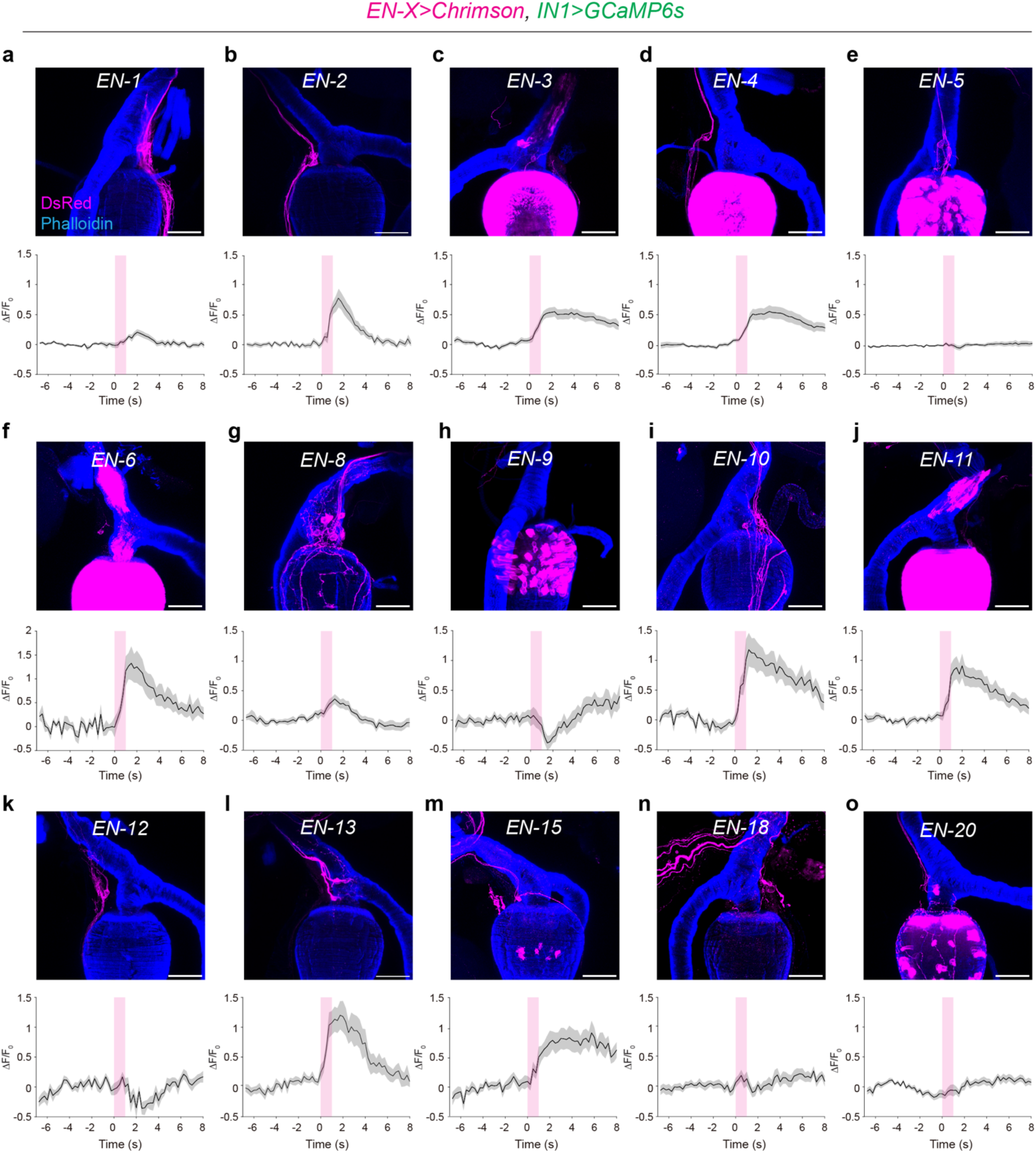
IN1s receive excitatory input from different classes of enteric neurons. **a-o,** Confocal images showing the expression patterns of EN split GAL4 lines expressing Chrimson (*ENs*> *Chrimson*) (magenta) in the HCG (top) (scale bars = 50μm). Normalised (ΔF/F_0_) GCaMP6s fluorescence in INs before and after optogenetic stimulation of different classes of enteric neurons (bottom) (n = 5-7 male flies, five trials per fly mean ± SEM).

**Extended Data Fig.5.**
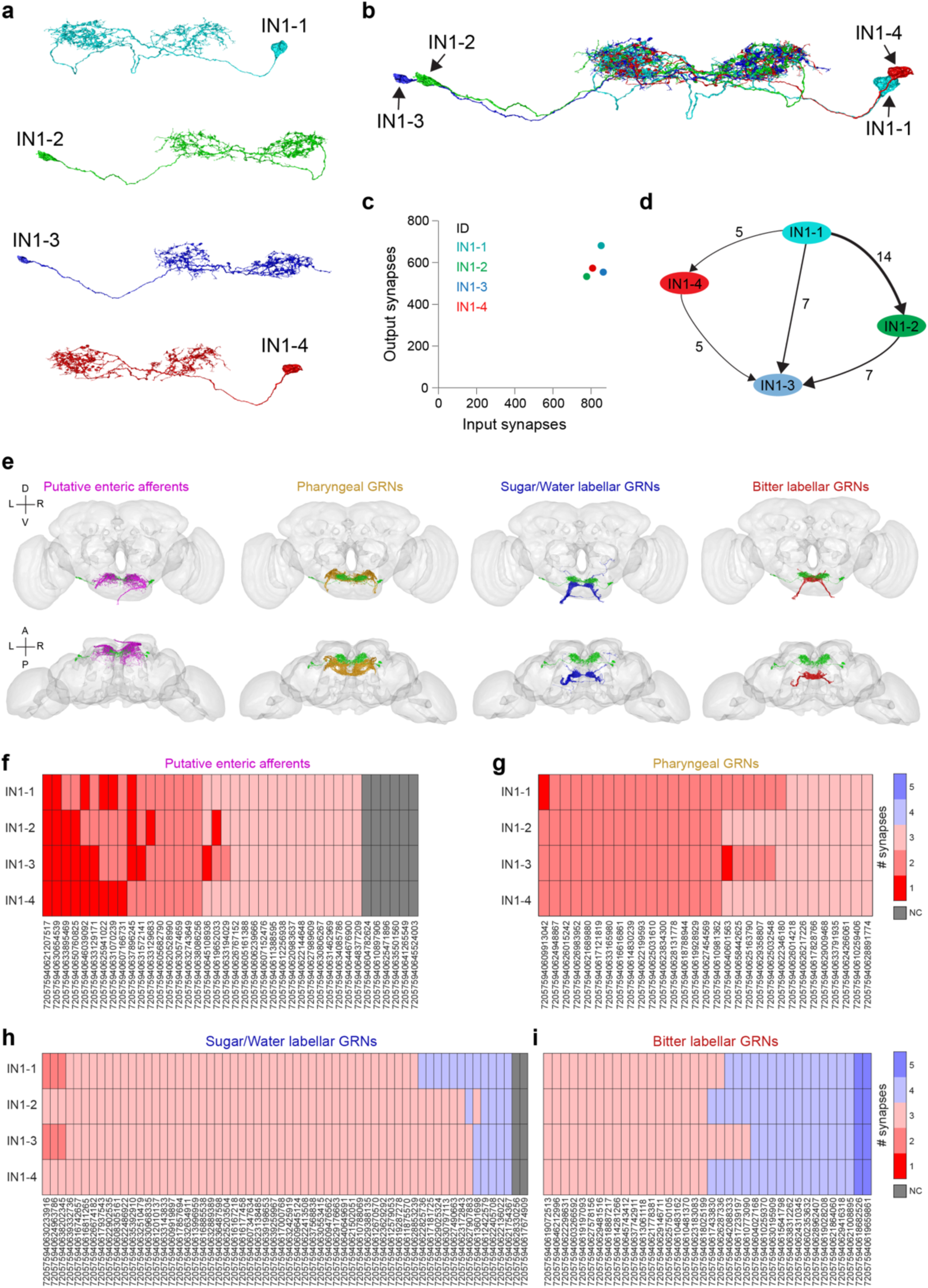
EM analysis of IN1s and their synaptic connectivity with different classes of GRNs and putative enteric neurons. **a-b**, EM reconstruction of putative IN1s (n=4) in the FAFB connectome is shown individually (**a**) or together (**b**). **c**, Total number of input and output synapses of putative IN1s in the FAFB connectome. **d**, Synaptic connectivity of IN1s to each other with arrows indicating the direction of connections (from presynaptic to postsynaptic neurons). The total number of synapses is shown on top of the arrows. **e**, EM reconstruction of putative IN1s (green) together with different classes of GRN (yellow, pharyngeal GRNs; blue, sweet/water labellar GRNs; red, bitter labellar GRNs) or putative enteric afferents (magenta) in the FAFB connectome. The front view (top) and top view (bottom) are shown. (L, left; R, right; D, dorsal; V, ventral; A, anterior; P, posterior). **f-i**, Heatmap showing the connectivity network between IN1s and different classes of GRNs or putative enteric afferents (NC, no path). IN1s are synaptically closer to putative enteric afferents than pharyngeal or labellar GRNs. A threshold of 5 synapses is used to determine connectivity between pairs of neurons.

**Extended Data Fig.6.**
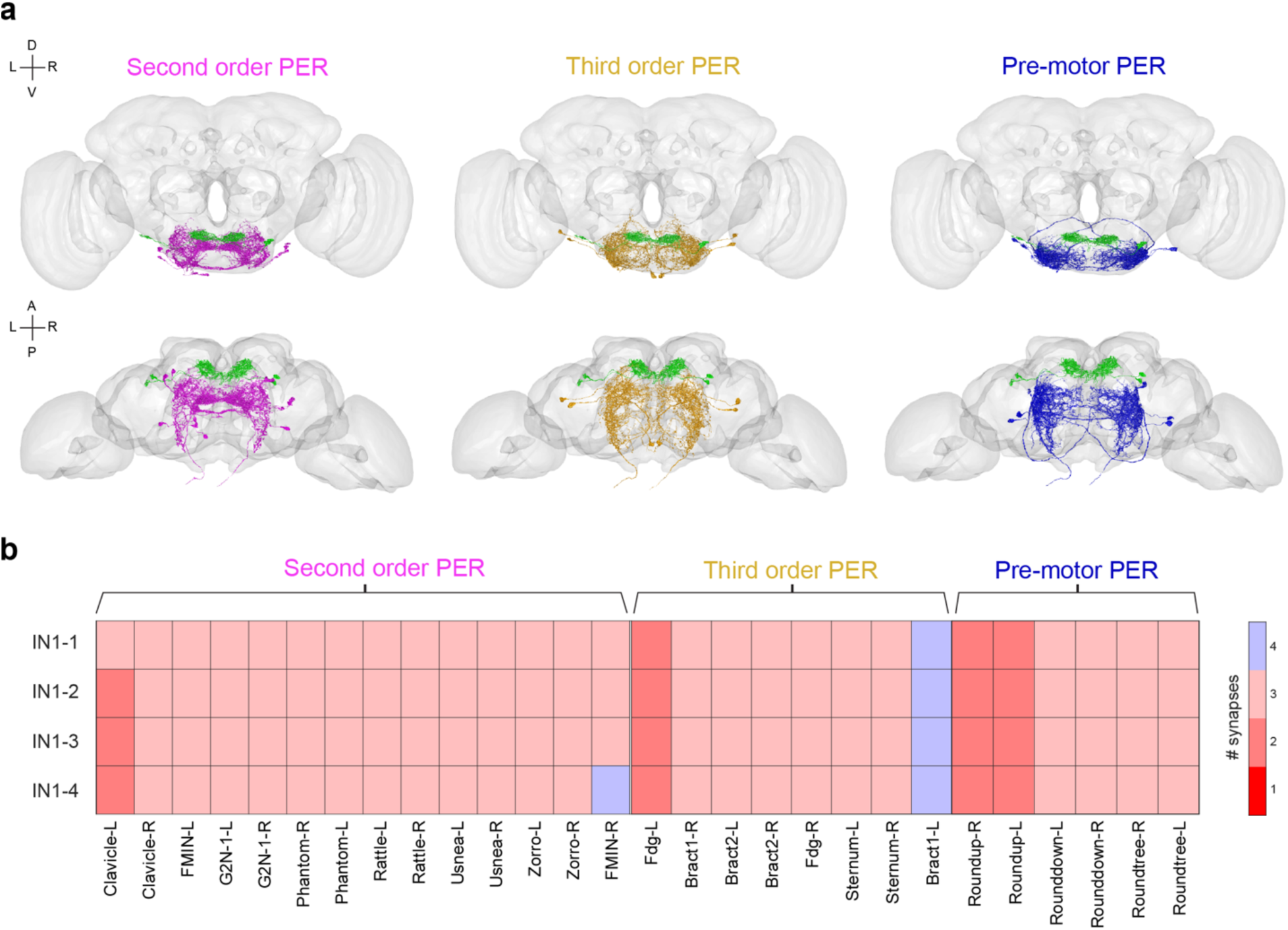
EM analysis of IN1s and their synaptic connectivity with different classes of PER neurons. **a**, EM reconstruction of putative IN1s (green) with different classes of PER neurons in the FAFB connectome (magenta, second order PER; yellow, third order PER; blue, pre-motor PER). The front view (top) and top view (bottom) are shown. (L, left; R, right; D, dorsal; V, ventral; A, anterior; P, posterior). **b**, Heatmap showing the connectivity network between IN1s and different classes of PER neurons. IN1s are at least two synapses away from PER neurons in the SEZ. A threshold of 5 synapses is used to determine connectivity between pairs of neurons. Neurons with cell bodies in the left hemisphere are labelled with -L, and neurons with cell bodies in the right hemisphere are labelled with -R.

**Extended Data Fig.7.**
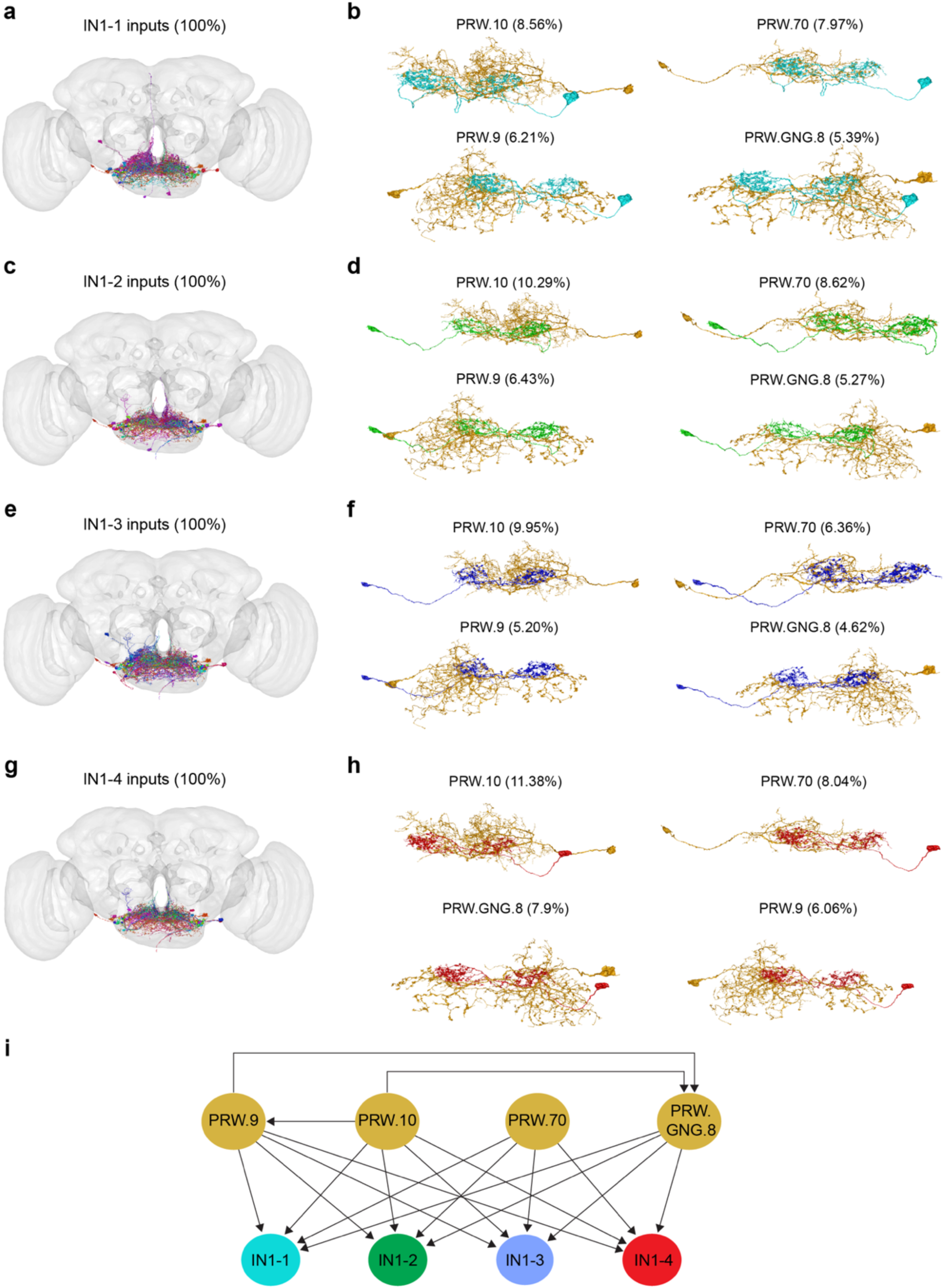
EM analysis of IN1 presynaptic neurons. **a-h**, EM reconstruction of putative IN1s (cyan, IN1-1; green, IN1-2; violet, IN1-3; red, IN1-4) with their presynaptic inputs (**a, c, e, g**). Anatomy of the top four presynaptic inputs to IN1-1 (**b**), IN1-2 (**d**), IN1-3 (**f**), or IN1-4 (**h**) in the FAFB brain dataset are shown. The input neurons are ordered based on their % synaptic input to IN1s. The Codex IDs of input neurons and the percentage of synaptic inputs they provide to IN1s are indicated on top of each panel. A threshold of five synapses is used to determine connectivity between neurons. **i**, Synaptic connectivity of IN1s and their inputs. Arrows indicate the direction of connections (from presynaptic to postsynaptic neurons). Synaptic connections between IN1s are not shown in this graph. Please see Extended Data Fig. 5d for recurrent connections between IN1s.

**Extended Data Fig.8.**
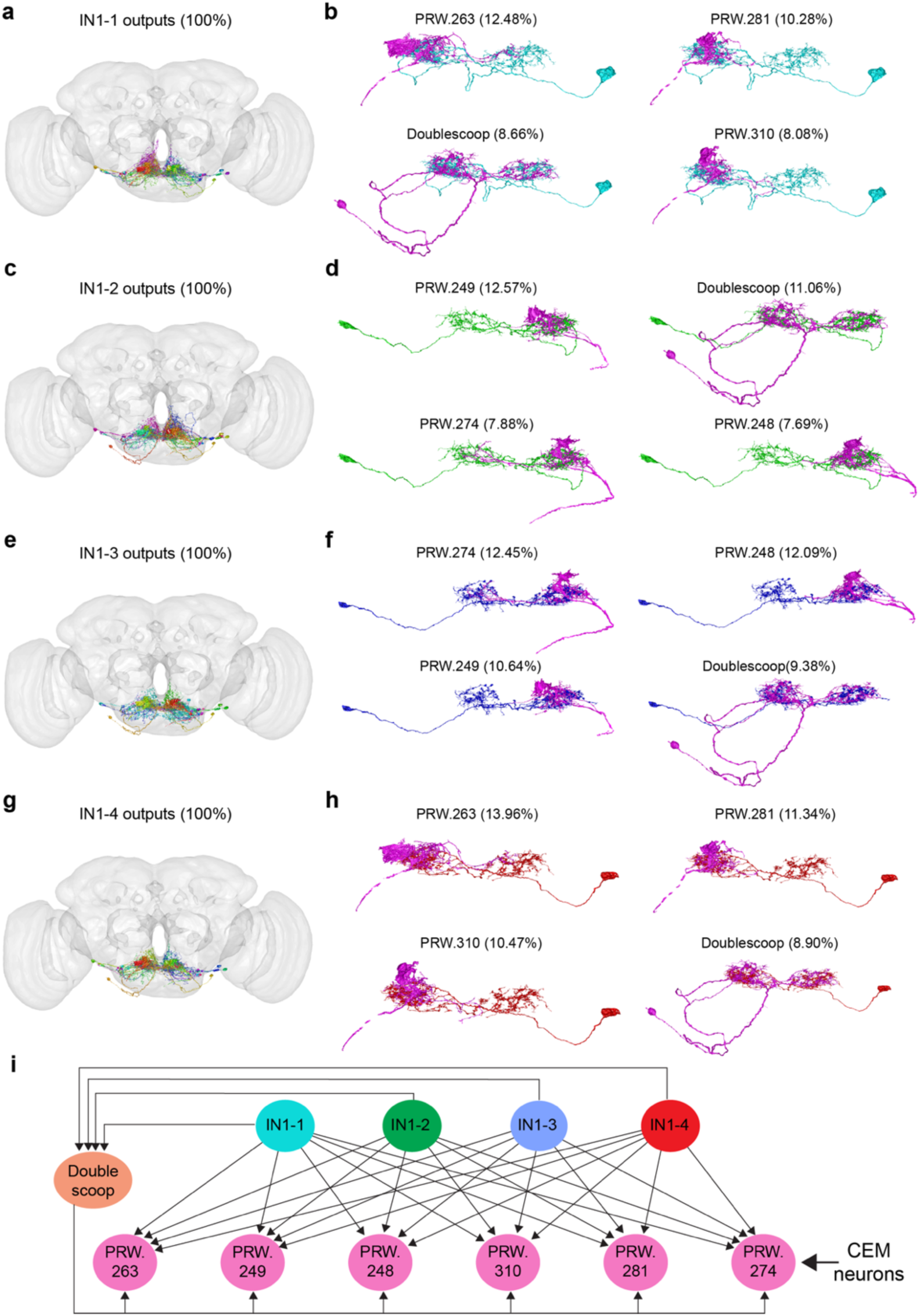
EM analysis of IN1 postynaptic neurons. **a-h**, EM reconstruction of putative IN1s (cyan, IN1-1; green, IN1-2; violet, IN1-3; red, IN1-4) with their postsynaptic outputs (**a, c, e, g**). Anatomy of the top four postsynaptic outputs for IN1-1 (**b**), IN1-2 (**d**), IN1-3 (**f**), or IN1-4 (**h**) in the FAFB brain dataset are shown. The Codex IDs of output neurons and the percentage of synaptic outputs IN1s provide to them are indicated on top of each panel. A threshold of five synapses is used to determine connectivity between neurons. **i**, Synaptic connectivity of IN1s and their outputs. Arrows indicate the direction of connections (from presynaptic to postsynaptic neurons). The six neurons (pink) shown in the bottom row (PRW.263, PRW.249, PRW.248, PRW.310, PRW.281, and PRW.274) are the crop-innervating enteric motor neurons (CEM neurons). Synaptic connections between IN1s are not shown in this graph. Please see Extended Data Fig. 5d for recurrent connections between IN1s.

**Extended Data Fig.9.**
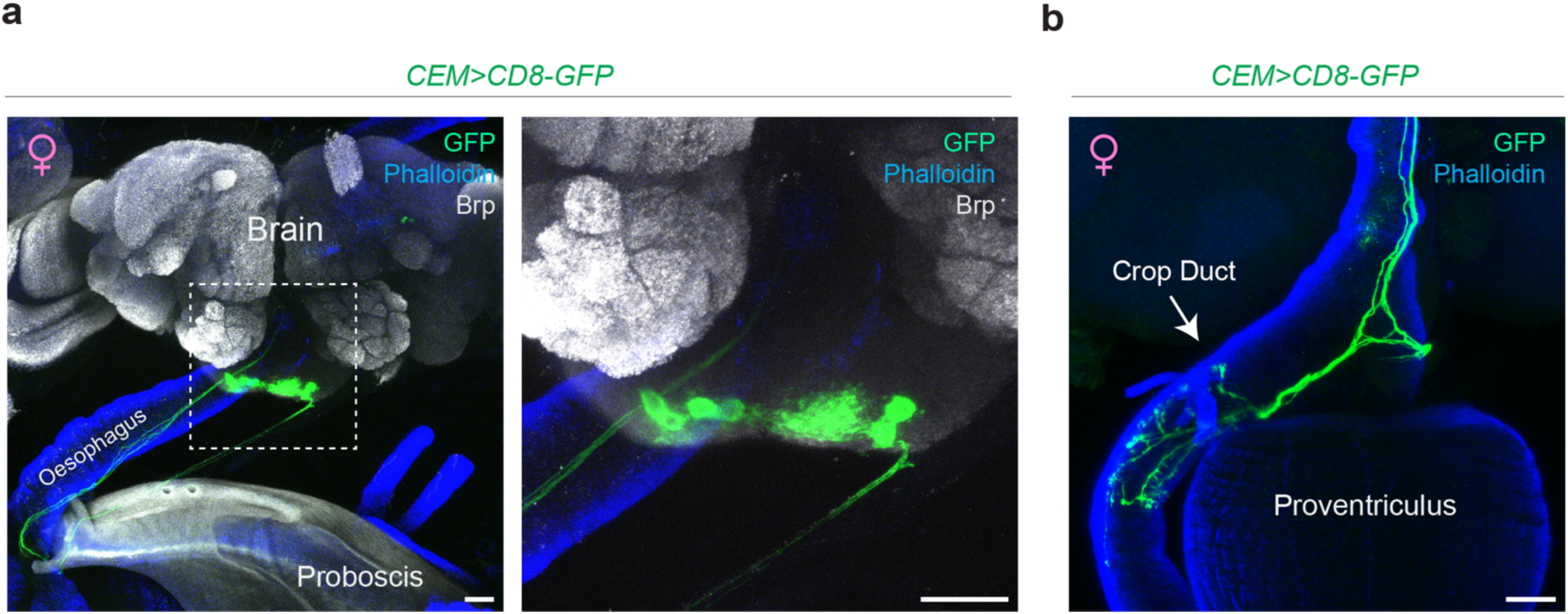
CEM neurons are present in both sexes. **a**, Confocal images of CEM neurons (green) in the female brain and their descending projections towards the oesophagus (blue) (scale bars = 25μm). **b,** Synaptic terminals of CEM neurons (green) innervating the crop duct (blue) in a female fly. (scale bar = 25μm).

**Extended Data Fig.10.**
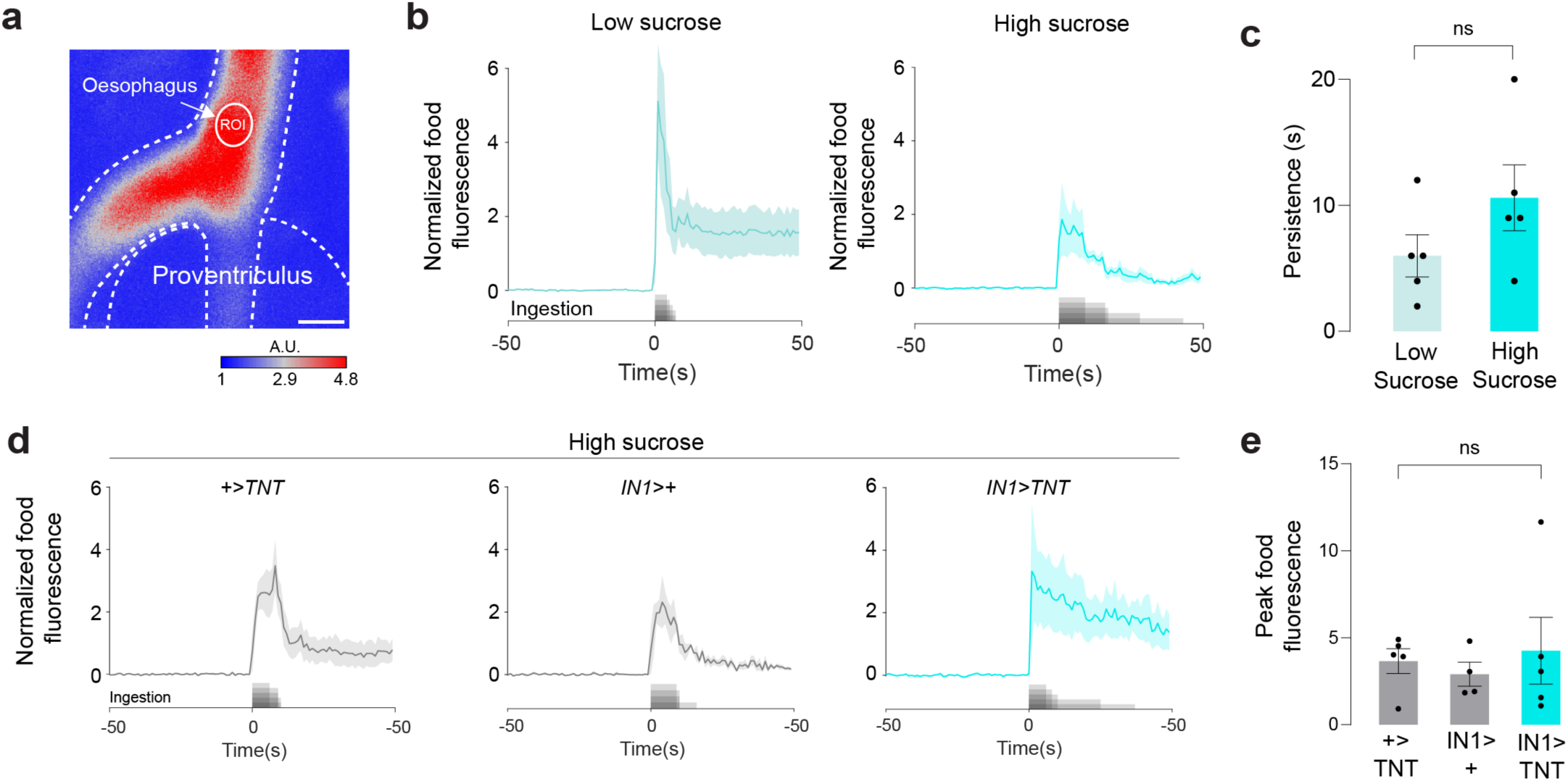
Inhibition of IN1s does not block entry of sucrose into the oesophagus. **a,** A representative image showing the ROI in the oesophagus during ingestion of fluorescein food (scale bar=25μm. A.U., arbitrary unit). **b**, Normalized food fluorescence in the oesophagus before and after ingestion of low (∼100mM) (left) and high (∼1M) (right) sucrose (n = 5 male flies, mean ± SEM). The sugar stimulus was provided *ad libitum*. **c**, Persistence of normalised food fluorescence in the oesophagus when flies are ingesting low (∼100mM) or high (∼1M) sucrose (n = 5 male flies, mean ± SEM, Unpaired t-test with Welch’s correction, p = 0.18). **d**, Normalized food fluorescence in the oesophagus before and after ingestion of high sucrose (∼1M) in flies with indicated genotypes (n = 4-5 flies per group; mean ± SEM). **e**, Peak food fluorescence in the oesophagus after ingestion of high sucrose (∼1M) in flies with indicated genotypes (n = 4-5 male flies; mean ± SEM). The amount of food entering the oesophagus is not altered when IN1 neurons are inhibited (n = 4-5 male flies, mean ± SEM, one-way ANOVA, p = 0.78).

## ONLINE METHODS

### Fly husbandry and genotypes

For all experiments, we used male flies 3 days post-eclosion unless otherwise noted. Flies were housed in a 25° C incubator with 60-65% humidity. Flies were grown on a conventional cornmeal-agar-molasses medium under a 12/12 light/dark cycle (lights on at 9 A.M.). When tested as controls, UAS or GAL4 stocks were tested as hemizygotes after crossing to w^1118^. IN1-split-GAL4 and IN1-split-LexA were generated in this study. IN1-split-GAL4 was generated by recombining 57F03-GAL4-DBD and 83F01-GAL4-AD on the 3^rd^ chromosome. IN1-split-LexA was generated by recombining 57F03-LexA-DBD and 83F01-GAL4-AD on the 3^rd^ chromosome. Gr43a^GAL4^ and Gr43a^LEXA^ were generously provided by Dr. Hubert Amrein^43^ (Texas A&M University). Ppk28-GAL4^35^ was generously provided by Dr. Micheal Gordon (The University of British Columbia). ChAT-Gal80^48^ was generously provided by Dr. Toshihiro Kitamoto (University of Iowa). Gr5a-GAL4^31^ and Gr66a-GAL4^31^ were generously provided by Dr. Kristin Scott (University of California, Berkeley). 10XUAS-Syn21-Chrimson88-tdT-3.1, LexAop2-Syn21-opGCaMP6s^27^ was generously provided by Dr. Michael Reiser (HHMI Janelia). 10xUAS-IVS-Syn21-Chrimson::tdT-3.1^25^ was generously provided by Dr. David Anderson (Caltech). LexAop-Chrimson-TdTomato^75^ was generously provided by Dr. John Tuthill (University of Washington). The following stocks were obtained from the BDSC: w^1118^ (5905); Gr43a-GAL4 (57637); Gr64f-GAL4 (57669); Gr64a-GAL4 (57661); Gr64d-GAL4 (57665); TMC-GAL4 (66557); Ir25a-GAL4 (41728); UAS-TNT-E (28837); UAS-CD8-GFP 3^rd^ chr. (32185); UAS-CD8-GFP 2^nd^ chr. (32186); UAS-CD8-RFP, LexAop-CD8-GFP (32229); UAS-GCaMP6s 2^nd^ chr (42746); UAS-GCaMP6s 3^rd^ chr (42749); UAS-GCaMP8s (92594); 15D05-GAL4 (48686); 17A11-GAL4 (48752); 20G03-GAL4 (48907); 24D12-GAL4 (49080); 25F11-GAL4 (49133); 37A08-GAL4 (49946); 38B05-GAL4 (49985); 44F09-GAL4 (50215); 70C02-GAL4 (39521); 15D05-Gla4.DBD (69218); R15D05-GAL4.AD (70556); R17A11-GAL4.DBD (68924); R20C05-GAL4.AD (70905); R20C10-GAL4.AD (70491); R20G03-GAL4.AD (70109); R20G03-GAL4.DBD (69047); R24D12-GAL4.AD (75677); R24D12-GAL4.DBD (68750); R25F11-GAL4.AD (70623); R25F11-GAL4.DBD (69578); R37A08-GAL4.AD (71028); R38B05-GAL4.DBD (69200); R44F09-GAL4.AD (71061); R70B03-GAL4.DBD (75656); R70C02-GAL4.DBD (69783); R84D10-GAL4.AD (70834). See Supplemental Table 1 for detailed information on fly genotypes in each figure.

### Transgenic fly production

57F03-LexA-DBD, 57F03-GAL4-DBD, and 83F01-GAL4-AD were generated in this study using Gateway recombination cloning. The 83F01 enhancer fragment was obtained from Dr. Gerry Rubin (HHMI Janelia) in a Gateway donor vector^64^. 57F03 enhancer is the same enhancer used to generate 57F03-GAL80 and 57F03-LexA^8^. We used the following Gateway destination vectors: ZpLexADBD_pBGUw (provided by Dr. Barry Dickson, Queensland Brain Institute), pBPZpGAL4DBDUw (Addgene plasmid #26233), pBPp65ADZpUw (Addgene plasmid #26234). The Transgenic fly lines were generated with the phiC31-based integration system^65^ (Best Gene Inc). The 57F03-GAL4-DBD and 57F03-LexA-DBD transgenes were inserted into the attP2 genomic locus, and the 83F01-GAL4-AD transgene was inserted into the VK00005 genomic locus.

### Immunohistochemistry and confocal microscopy

All brains, ventral nerve cords, and guts were dissected in 1x phosphate-buffered saline (PBS, diluted from 10 × PBS listed in the resource table). The samples were then stained as previously described^8^. Briefly, immediately after dissection, samples were transferred to a 1.5mL centrifuge tube filled with ∼200ul of 1xPBS using a pipette. After all sample collection was completed, the 1xPBS was removed from the centrifuge tube, and samples were incubated in 4% paraformaldehyde (PFA, Electron Microscopy Sciences, Cat# 15711) on an orbital shaker for 15 to 25 minutes. After tissue fixation, samples were washed with PBT for 4×15 minutes. For most immunohistochemistry experiments in this study, we used 0.1% PBT. However, for the experiment involving the VGLUT antibody, we used 0.2% PBT. Next, the samples were incubated with 5% Normal Goat Serum diluted in PBT (NGS-PBT) for approximately 30 minutes, followed by incubation with primary antibodies diluted in NGS-PBT for approximately 2-5 days at 4°C. After the primary antibody incubation, samples were washed with PBT for 5×15 minutes and incubated with secondary antibodies diluted in NGS-PBT for ∼24 to 48 hours at 4°C. Once the antibody incubations were completed, samples were washed with PBT at room temperature for 4×15 minutes and incubated with Slowfade medium (ThermoFisher Scientific, Cat. # S36936) on an orbital shaker for ∼30 minutes before getting mounted on a microscope slide. The samples were covered by a glass coverslip and sealed using clear nail polish (Clear Nail Polish, Electron Microscopy Sciences, Cat. # 72180). The following primary and secondary antibodies were used: chicken polyclonal anti-GFP, 1:3000 (Abcam Cat# ab13970), rabbit polyclonal anti-DsRed, 1:500 (Takara Bio, Cat# 632496), mouse monoclonal anti-Bruchpilot, 1:20 (DSHB, Cat# Nc82), rabbit anti-VGLUT, 1:500 (kindly provided by Dr. Dion Dickman), rat monoclonal anti-elav, 1:30 (DSHB, Cat# Rat-Elav-7E8A10), Phalloidin (Sigma, Cat#P2141), Alexa 546-conjugated goat anti-rabbit IgG, 1:1000 (Invitrogen, Cat# A11035), Cyanine5-conjugated goat anti-mouse IgG, 1:500 (Invitrogen, Cat# A10524), Alexa 633-conjugated goat anti-mouse IgG, 1:500 (Invitrogen, Cat# A-21052), Alexa 488-conjugated goat anti-chicken IgG, 1:1000 (Invitrogen, Cat# A11039). Samples were mounted with Slowfade medium (ThermoFisher Scientific, Cat. # S36936). For Extended Data Figs. 2a-b, samples were collected and immediately embedded in an imaging medium (Tissue-Tek® O.C.T. Compound, Sakura) and sealed using a coverslip. All fluorescent images were taken using a Zeiss LSM880 upright confocal microscope and Zeiss digital image processing software ZEN. Z-stacks were acquired at 1024×1024-pixel resolution with a z-step size of 1 to 5 μm.

### Fly preparation before two-photon imaging

For two-photon imaging coupled with optogenetics experiments, two-day-old male flies were fasted with or without ATR (All-*trans*-retinal, Sigma-Aldrich Cat#R2500, concentration = 0.5mM) in a vial containing Kimwipe soaked in 1ml MilliQ water. An aluminium foil was wrapped around the vial to protect the ATR from light exposure. The fasting duration and/or ATR treatment lasted 18 to 26 hours right before the imaging experiment. For two-photon imaging during food ingestion experiments, two-day-old male flies were fasted for 18-26 hours in a vial containing Kimwipe soaked in 1ml MilliQ water. To generate flies that are in a fed state, we transferred the previously fasted flies into a vial with ∼1M sucrose with 0.02% (m/v, or 2g/l) brilliant blue (Sigma-Aldrich, Cat# 80717) dye 1-4 hours before the imaging experiment. Flies that showed a blue colour in their abdomen were used in the two-photon imaging.

### Fly mounting and dissection for calcium imaging in the brain

Flies were prepared as previously described^8^. A custom-made fly holder was used for all two-photon *in vivo* imaging experiments. On the day of the experiment, a male fly was anaesthetised briefly with CO_2_ and tethered to a piece of transparent tape (Scotch® Transparent Tape) covering the hole in the fly holder. The fly head was secured using a human hair placed across the fly neck. We removed the tibia and tarsal segments of the forelegs during imaging experiments that involved food delivery. For the optogenetic imaging experiments, the proboscis of the fly was fully extended by fine forceps (Dumont #5, FST, Cat#11254-20) and fixed using UV curable adhesive (Bondic®) in a fully extended position to minimise the movement during imaging. Next, a small hole was cut into the tape, precisely above the head, to allow the head capsule to extend above the plane of the tape. UV curable adhesive was applied to the fly’s eyes and anterior and posterior parts of the head to restrict head movement. Once the fly’s head was fixed, ∼0.35ml of adult hemolymph-like (AHL) saline (108mM NaCl, 5mM KCl, 8.2mM MgCl_2_·6H_2_O, 2mM CaCl_2_·2H_2_O, 4mM NaHCO_3_, 1mM NaH_2_PO_4_·2H_2_O, 5mM Trehalose·2H_2_O, 10mM Sucrose, 5mM HEPES, pH adjusted to 7.5) was applied on top, and the head capsule was opened by carefully cutting the cuticle covering the dorsal-anterior porting of the fly head, including the antennae. Finally, we removed the obstructing air sacks and fat bodies using fine forceps to gain better optical access to the fly brain. The fly holder was then placed under the two-photon microscope for imaging.

### Fly mounting and dissection for calcium imaging in the gastrointestinal tract

On the day of the experiment, a male fly was anaesthetised briefly with CO_2_ and tethered to a piece of transparent tape (Scotch® Transparent Tape), covering the hole in the fly holder. The fly head and body were secured using human hair, one placed across the fly neck, the other onto the abdomen segment, between the midlegs and the hindlegs. The tibia and tarsal segments of the forelegs were then removed to avoid disruption of food delivery during imaging. Similar to brain imaging preparation, a small hole was cut into the tape, precisely above the head plus the thorax segment, to allow the head and the thorax segment to extend above the plane of the tape. UV curable adhesive (Bondic®) was then applied to seal the space between the fly’s body and the transparent tape. We checked the ability of flies to extend their proboscis after the fixation to ensure they can ingest food during imaging. Once the fly’s head and thorax were fixed ∼0.35ml of AHL saline (108mM NaCl, 5mM KCl, 8.2mM MgCl_2_·6H_2_O, 2mM CaCl_2_·2H_2_O, 4mM NaHCO_3_, 1mM NaH_2_PO_4_·2H_2_O, 5mM Trehalose·2H_2_O, 10mM Sucrose, 5mM HEPES, pH adjusted to 7.5) was applied on top of the thorax and the thorax cuticle, muscles, air sacks and fat bodies covering the hypocerebral ganglion were removed to gain optical access to the enteric neurons. The fly holder was then placed under the two-photon microscope for imaging.

### Two-photon imaging

All functional imaging experiments were performed using a resonant scanning two-photon microscope (Bergamo II, Thorlabs) equipped with a 16X Plan Fluor Objective (Nikon, N16XLWD-PF) and GaAsP detectors (Hamamatsu). We used ThorImage software (Thorlabs, v4.0.2020.2171) to control the microscope. Two-photon excitation was provided by a Chameleon Ti: Sapphire femtosecond pulsed laser with pre-compensation (Vision II, Coherent) centred at 920 nm. The laser was directed through a resonant scanning galvanometer for fast-scanning volumetric imaging, and a piezo-electric Z-focus controlled the objective. Laser power was measured using a power meter (PM100D with S175C, Thorlabs). Laser power after the objective ranged between ∼25-35 mW for brain imaging and ∼10-60 mW for enteric imaging. Before the functional imaging trials, we took a whole-brain z-stack to ensure Chrimson-tomato and/or GCaMP6s proteins were adequately expressed in the brain or the enteric neurons. We then focused on the selected region of interest (ROI) and recorded 4-minute (for all trials with optogenetics stimulation) or 8-minute (for all trials without optogenetics stimulation) volumetric time-lapse GCaMP6s, GCaMP8s or fluorescein fluorescence. The starting and ending z-position of the volumetric imaging is determined to cover the whole region of interest. The details of the fast volumetric scanning can be found in the table below:

| ROI | z-planes | Scan rate (Hz) | Step size (µm) | Resolution (pixel) | Figure |
| --- | --- | --- | --- | --- | --- |
| IN1 projections | 8 | 4.63 | 10 | 256×256 | Fig. 1,2, Extended Data Fig. 4 |
| Gr43a cell bodies | 8-10 | 3.95-4.63 | 10 | 256×256 | Fig. 3 |
| EN cell bodies | 10 | 3.95 | 10 | 256×256 | Fig. 4 |
| CEM projections | 8-10 | 3.95-4.56 | 10-15 | 256×256 | Fig. 5 |
| Fluorescent food ingestion | 10 | 2.02-2.03 | 20 | 512×512 | Fig. 6, Extended Data Fig. 10 |

We used an infrared light (JC Infrared Illuminator) and a FLIR Blackfly-S (BFS-U3-16S2M) equipped with a zoom lens (MLM3X-MP, 0.3X-1X, 1:4.5; Computar) and a Near-Infrared bandpass filter (BP810-34, Midwest Optical Systems, INC.) inside the imaging chamber to record the motion of the fly during imaging experiments (Software: SpinView, Spinnaker v. 2.0.0.147, FLIR Systems, Inc.). The video (30 fps) and the two-photon imaging data acquired by ThorImage were synchronised using GPIO connections. The ThorImage and LED optogenetic stimulation were synchronised using ThorSync software (Thorlabs, version 4.1.2020.1131). In the optogenetics calcium imaging trials that do not involve food ingestion, a spherical treadmill supported by an air pump was placed below the fly to minimise stress during imaging. In the imaging trials that involved food ingestion, the ball was removed to generate space for the food delivery device. At the end of each imaging experiment, we assessed the flies’ health condition by mechanical stimulation of the leg using forceps. The data collected from flies that did not respond to the mechanical stimulus were excluded from the final data analysis because we considered those flies unhealthy. Furthermore, imaging data from flies that showed substantial movement in the Z-direction were also discarded on rare occasions because of the severe motion artefacts in the calcium trace.

### Optogenetic stimulation during two-photon imaging

Optogenetic stimulation was generated using a 617nm LED, which is integrated into the light path of the two-photon microscope and delivered to the fly brain via the objective. LED light intensity (∼0.75mW) was measured after the objective by an optical power meter (PM100D, Thorlabs) equipped with a light intensity sensor (S175, Thorlabs). A long-pass filter (FELH0600, Thorlabs) was used to reduce the background elevation caused by 617nm-LED light during optogenetic stimulation. All optogenetic activation experiments started with ∼30s scanning without stimulation to capture baseline GCaMP fluorescence. Next, LED light stimulation was continuously delivered to the fly brain for 1s or 10s. The stimulation was repeated five times at 30s intervals. For the optogenetic stimulation during fluorescent food ingestion (Fig. 6f-h), a continuous 30s long optogenetic stimulation was applied at around t=31-61s of the four-minute two-photon imaging trial.

### Sugar ingestion in tethered flies

The sucrose solution was prepared by dissolving sucrose (Sigma-Aldrich, Cat# 9378) in MilliQ water. For high-concentration sucrose solution (∼1M), 0.34g sucrose was dissolved in 1 ml MilliQ water. For low-concentration sucrose solution food (∼100mM), 0.017g sucrose was dissolved in 500µl MilliQ water. In all ingestion experiments with two-photon imaging, except for fluorescent food ingestion, Brilliant Blue (2 g/l) (Sigma-Aldrich, Cat# 80717) was added to the high- or low-concentration sucrose solution. This allowed us to confirm ingestion episodes by inspecting the blue dye presence in the fly gut after the experiments. In fluorescent food ingestion imaging experiments, fluorescein (0.5 g/l) (Dextran Fluorescein, Thermo Fisher, Cat# D1823) was added to the high- or low-concentration sucrose solution. The sucrose solution was presented to the fly using a pulled glass capillary attached to a microinjector (Drummond Nanoject II Auto, CAT # 3-000-204) to deliver the sucrose solution in precise volumes. To prevent the sucrose solution from wicking down the sides of the capillary, we applied dental wax to the exterior of the glass capillary. We used a micromanipulator to control the Nanojet and the capillary’s movement during imaging (Siskiyou, Micromanipulator Controller Mc1000e). The sugar stimulation was present in discrete durations during imaging experiments unless otherwise stated.

### Data processing and analysis

#### Confocal image processing

Confocal images were processed using the FIJI open-source image-processing package (https://imagej.net/software/fiji/) or Imaris (Oxford Instruments, Imaris x64, version 9.9.0). All confocal images shown in this paper, except for the cross-section images in Figs. 3i, Fig. 4f, and Extended Data Fig. 3k, are z-projections of the confocal image stacks. Confocal image stacks of the fly gastrointestinal tract in Figs. 3i, Fig. 4f, and Extended Data Fig. 3k were processed using Imaris to generate the cross-section images in X, Y, and Z directions.

#### Two-photon functional imaging data processing

All two-photon imaging data processing was completed using custom-made code written in (version R2022b). Two-photon volumetric image frames were projected along the z-axis for each trial and then aligned by translating each frame in the x and y plane using the MATLAB register function. Registration results were manually inspected to avoid the artefacts produced by the movement of ROI as much as possible. If the registration result from MATLAB were not ideal, image stacks were registered for the second round using TurboReg (https://bigwww.epfl.ch/thevenaz/turboreg) or manually using FIJI (https://imagej.net/software/fiji/). Image stacks that failed the registration process were discarded. Region of interest (ROI) selection was achieved by manually drawing one or multiple mask(s) surrounding the cell bodies or the neuronal projections using MATLAB freehand function. The ROI masks were applied to all z-projected frames, and the average grey value within each ROI was extracted from each frame to generate a time series.

| Two photon imaging | ROI | Figure |
| --- | --- | --- |
| IN1 | Projections in left or right hemisphere | Fig. 1,2, Extended Data Fig. 4 |
| Gr43a | Cell bodies | Fig. 3 |
| ENs | Cell bodies | Fig. 4 |
| CEM | Projections in the gut | Fig. 5 |
| Fluorescent food ingestion | Crop duct or oesophagus | Fig. 6, Extended Data Fig. 10 |

##### Optogenetic stimulation trials

(Fig. 1c-g, Fig.2 a-f, Fig. 5d-e, Extended Data Fig. 4a-o): During optogenetic stimulation, the LED light-induced a slight background elevation observable in the imaging data. This background elevation served as an indicator of stimulation ON and OFF times during data processing. Background subtraction was applied to each imaging frame during data processing to eliminate this noise. To calculate ΔF/F_0_ during optogenetic stimulation trials, the fluorescent time series was first chopped into five segments corresponding to the five stimulation episodes. For 1s-stimulation trials, each imaging segment consists of the time +/-7s pre- and post-stimulation. For 10s-stimulation trials, each imaging segment consists of the time +/- 10s pre- and post-stimulation. To calculate the ΔF/F_0_, we first subtracted the background fluorescence value from the ROI fluorescence value for each frame. Next, the baseline fluorescence F_0_ was calculated by averaging the fluorescent intensities from 5s (t=-6 to -1 seconds) before the stimulation onset (t=0). Finally, ΔF/F_0_ was calculated using the following formula: 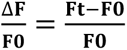. (**F0**=fluorescence at baseline, **Ft**=fluorescence at time t). The resulting time series was binned into 0.25s time bins and plotted as ± SEM ΔF/F_0_. We averaged the ΔF/F_0_ across all trials to calculate the average and SEM.

##### Sucrose ingestion trials

(Fig. 3d, e, Fig. 4d, Fig. 5f, Fig. 6b, d): To calculate ΔF/F_0_ during sucrose ingestion trials, ROI fluorescent intensities recorded from +/- 50s pre- and post-ingestion were used unless otherwise stated. Background subtraction was not applied to the fluorescent time series data in these trials. Baseline fluorescence F_0_ was calculated by averaging the fluorescent intensities from 10s (t=-10s to 0s) before the ingestion onset. ΔF/F_0_ was calculated using the same formula as in the optogenetic stimulation experiments. For EN cell body imaging (Fig. 4d), if an imaging trial contained multiple ROIs corresponding to multiple cell bodies, to generate the averaged ΔF/F_0_ plot, ΔF/F_0_ traces from all ROIs were first averaged within a fly and then averaged across flies. For Gr43a cell body imaging (Fig. 3d, e), ROI fluorescent intensities recorded from +/- 30s pre- and post-ingestion were used. In these imaging experiments, we noticed not all Gr43a neurons responded to sucrose ingestion. Only neurons that were activated by sucrose were used in the data analysis. A Gr43a neuron was classified as responsive if the peak ΔF/F_0_ signal was 3x greater than the ΔF/F_0_ standard deviation above baseline.

##### Optogenetic stimulation and ingestion

(Fig. 6g-j): For combined imaging trials with optogenetic stimulation and ingestion, ROI fluorescent intensities recorded from +/- 5s pre- and post-ingestion were used. ΔF/F_0_ was calculated following the same steps as for optogenetic stimulation imaging trials, with the F_0_ time window set from 3 seconds (t=- 3 to 0 seconds) before ingestion onset. Background subtraction was applied to each imaging frame during processing.

##### Peak ΔF/F_0_ (or peak food fluorescence) calculations

Throughout this study, peak ΔF/F_0_ is defined as the maximum or minimum value of ΔF/F_0_ within a specified time window. Time windows for peak ΔF/F_0_ (or peak food fluorescence) calculations are as follows: in Fig. 1h and Fig. 4b, 0 to 4 seconds after optogenetic stimulation onset; in Fig. 3f, the duration of the ingestion bout; in Fig. 5g, 0 to 50 seconds after ingestion onset; in Fig. 6e and Extended Data Fig. 10e, the duration of the ingestion bout; in Fig. 6h, j, 0 to 5 seconds after ingestion onset. We plotted the average peak ΔF/F_0_ (or peak food fluorescence) and used GraphPad Prism for statistical comparisons.

##### Persistence calculations

Persistence in Fig. 3g, Fig. 6c, and Extended Data Fig. 10c is the total duration at half maximum (FDHM), calculated as the duration between the points where the peak ΔF/F_0_ is half its maximum value. Persistence in Fig. 5h is calculated as the total duration where the ΔF/F_0_ is lower than half its minimum value after ingestion onset (t=0-10s). We plotted the average peak ΔF/F_0_ (or peak food fluorescence) and ΔF/F_0_ persistence and used GraphPad Prism for statistical comparisons. We plotted the average persistence and used GraphPad Prism for statistical comparisons.

### Data exclusion

Flies that appeared in poor health after imaging and/or had low-quality image data due to motion artefacts or other reasons were excluded from data analysis. These were less than 4% of the flies used in the entire study.

### EM analysis

We used the Flywire open-source platform to identify and classify the putative IN1 neurons^52^. We first identified putative IN1s based on light microscopy data, projection patterns, and cell body locations. To plot the putative IN1s in a standard brain mesh, we used the natverse package in R-studio (version R4.3.3), a toolbox for combining and analysing neuroanatomical data^76^. Natverse consists of multiple R-packages that allow the analysis of light microscopy and EM datasets across various model organisms, including *Drosophila melanogaster*. We mainly used the R “fafbseg” package to access and analyse the Flywire datasets. Details of the “fafbseg” package can be found at https://natverse.org/fafbseg/. First, we downloaded neuron meshes for each putative IN1 neuron from Flywire into the R environment and visualised them in 3D using the FAFB14 standard brain mesh. Next, we used Flywire to automatically detect synaptic sites across putative IN1 neurons (n=4) and generated the connectivity matrix across IN1 neurons using Codex (Connectome Data Explorer: codex.flywire.ai)^60^. We also used Codex to identify the input and output neurons of IN1s. The neuron meshes of the IN1 outputs and inputs, as well as the candidate IN1 interacting neurons, such as sugar, bitter GRNs, second and third-order taste neurons and taste motor neurons, were downloaded from Flywire into the R environment and visualised in 3D using the FAFB14 standard brain mesh. The connectivity heatmaps for these neurons were generated from the data in Pathway analysis in Codex using a costume MATLAB script. The MATLAB script is available at https://github.com/Nilayyapici/Cui_et_al. The neuron IDs used in this paper are listed in Supplementary Table 2.

### Statistical tests

Sample sizes used in this study were based on previous literature in the field. Experimenters were not blinded in most conditions, as almost all data analysis were automated and done using a standardised computer code. All statistical analysis were performed using Prism Software (GraphPad, Version 10.1.1). One-way ANOVA with Bonferroni post hoc multiple comparisons (for data that are mainly normally distributed) or non-parametric Kruskal-Wallis test with Dunn’s post hoc multiple comparisons (for data that are not normally distributed) were used to compare more than two genotypes or conditions (for details, see the legends of each figure). The paired or unpaired t-test (two-tailed) with Welch’s correction was used to compare two genotypes or conditions. Data labelled with different letters are statistically different.

